# Proteomic traits vary across taxa in a coastal Antarctic phytoplankton bloom

**DOI:** 10.1101/2021.05.05.442818

**Authors:** J. Scott P. McCain, Andrew E. Allen, Erin M. Bertrand

## Abstract

Production and use of proteins is under strong selection in microbes, but it’s unclear how proteome-level traits relate to ecological strategies. We identified and quantified proteomic traits of eukaryotic and prokaryotic microbes through an Antarctic phytoplankton bloom using in situ metaproteomics. To do this, we first used simulations, cultures, and bioinformatic methods to rigorously assess our inferences about various proteomic traits and use these assessments to provide several practical recommendations for researchers using metaproteomics. Different taxa, rather than different environmental conditions, formed distinct clusters based on their ribosomal and photosynthetic proteomic proportions, and we propose that these characteristics relate to ecological differences. We defined and used a proteomic proxy for regulatory cost, which showed that SAR11 had the lowest regulatory cost of any taxa we observed at our summertime Southern Ocean study site. Haptophytes had lower regulatory cost than diatoms, which may underpin haptophyte-to-diatom bloom progression in the Ross Sea. Using metaproteomics, we have quantified several proteomic traits (ribosomal and photosynthetic proteomic proportions, regulatory cost) in eukaryotic and prokaryotic taxa, which can then be incorporated into trait-based models of microbial communities that reflect resource allocation strategies.

## 1 Introduction

Microbes are constantly faced with an optimization problem: which proteins should be produced, when, and how many? The solutions to this problem dictate metabolic rates, cell stoichiometry, and taxonomic distribution [1, 2, 3, 4, 5]. Yet, it’s unclear what these solutions actually are in terms of proteome composition, and if different microbes have arrived at different solutions. Microbes are typically compared based on their unique repertoires of potential proteins [e.g. 6, 7, 8], but taxa have shared proteins as well – are these shared proteins produced in similar amounts? Or, do taxa produce distinct amounts under identical conditions? Diverse taxa produce proteins in strikingly similar ratios within some pathways [9], but is stoichiometry conserved between pathways? The answers to these questions will direct future efforts for modelling microbial communities. Perhaps microbes can be represented as collections of genes [10, 11], or, perhaps variation in proteome composition will shed light on the underpinnings of their ecological strategies and biogeochemical contributions.

Ecological strategies must be tied to gene expression [12], and models can experimentally test hypotheses to evaluate such connections. Material models (i.e. cultures) have clearly demonstrated that selection acts strongly on protein production [13, 14, 15]. While powerful, these approaches are limited to only a few culturable organisms, which can overlook core differences found in less-studied organisms [e.g. 16]. Computational models have also characterized trade-offs and metabolic behaviours in microbes [e.g. 17, 18, 19]. While models are critical from a reductionist perspective, characterization and prediction of microbial activity in their environments remains a central research goal.

Observing and measuring resource allocation in microbes *in situ* can also link gene expression to ecological strategies [e.g. 5, 20, 21, 22, 23, 24]. For example, diatom and haptophyte transcriptional dynamics reflect their distinct growth strategies, assessed using metatranscriptomics [22, 23]. Metaproteomics has similarly identified increased abundance of transporter proteins across an oceanographic gradient of decreasing nutrients [5]. Both of these meta-omic approaches can quantify *in situ* resource allocation, but proteins cost more to produce and therefore better reflect resource allocation [25]. To our knowledge, metaproteomics has not been used to quantify variation in resource allocation strategies across eukaryotic and prokaryotic microbes.

Our objective was to identify and quantify proteomic ‘traits’ for various eukaryotes and prokaryotes, by examining microbial proteome composition through a four-week time series at the Antarctic sea ice edge. We define a proteomic trait as a characteristic of an organism at the proteome-level, that includes both the abundance and identity of a protein (or group of proteins), and is connected to organismal fitness or performance [26]. Metaproteomics is confronted by several methodological issues and biases, which we rigorously assess. We subsequently provide practical recommendations for researchers using metaproteomics to examine microbial resource allocation. Next we connect proteomic resource allocation to the ecology of these plankton. Lastly, we suggest that characterizing coarse-grained proteomes, which we define as a grouping of functionally related proteins, may be useful for assessing nutrient deficiency in the ocean.

## 2 Methods

### 2.1 Field sampling

We collected samples once per week over four weeks at the Antarctic sea ice edge, in McMurdo Sound, Antarctica [December 28, 2014 ‘GOS-927’; January 6 ‘GOS-930’, 15 ‘GOS-933’, and 22 ‘GOS-935’, 2015; as previously described in 27]. Sea water (150–250 L) was pumped sequentially through three filters of decreasing size (3.0, 0.8, and 0.1 um, 293 mm Supor filters). Separate filter sets were acquired for metagenomic, metatranscriptomic, and metaproteomic analyses, over the course of ~3 h, each week (36 filters in total). Filters for nucleic acid analyses were preserved with a sucrose-based buffer (20 mM EDTA, 400 mM NaCl, 0.75 M sucrose, 50 mM Tris-HCl, pH 8.0) with RNAlater (Life Technologies, Inc.) for the nucleic acid filters and without RNAlater for the protein filter, flash frozen in liquid nitrogen in the field and subsequently stored at −80 C until processed in the laboratory.

### 2.2 Metagenomic and metatranscriptomic sequencing

We used metagenomics and metatranscriptomics to obtain reference databases of potential proteins for metaproteomics. We additionally used a database assembled from a similarly processed metatranscriptomic incubation experiment [28], conducted with source water from the January 15, 2015 time point (these samples were collected on a 0.2um Sterivex filter and processed as previously described).

For samples from the GOS-927, GOS-930, GOS-933 and GOS-935 filters, RNA was purified from a DNA and RNA mixture [29]. 2 *μ*g of the DNA and RNA mixture was treated with 1 *μ*l of DNase (2 *μ*/*μ*l; Turbo DNase, TURBO DNase, ThermoFisher Scientific), followed by processing with an RNA Clean and Concentrator kit (Zymo Research). An Agilent TapeStation 2200 was used to observe and verify the quality of RNA. 200 ng of total RNA was used as input for rRNA removal using Ribo-Zero (Illumina) with a mixture of plant, bacterial, and human/mouse/rat Removal Solution in a ratio of 2:1:1. An Agilent TapeStation 2200 was used to subsequently observe and verify the quality of rRNA removal from total RNA. rRNA-deplete total RNA was used for cDNA synthesis with the Ovation RNA-Seq System V2 (TECAN, Redwood City, USA). DNA was extracted for metagenomics from the field samples (GOS-927, GOS-930, GOS-933 and GOS-935) according to [29]. RNase digestion was performed with 10 *μ*l of RNase A (20 mg/ml) and 6.8 ul of RNase T1 (1000U/*μ*l), which were added to 2 *μ*g of genomic DNA and RNA mixture in a total volume of 100 *μ*l, followed by 1 hour incubation at 37°C and subsequent ethanol precipitation in −20°C overnight.

Samples of double stranded cDNA and DNA were fragmented using a Covaries E210 system with the target size of 400 bp. 100 ng of fragmented cDNA or DNA was used as input into the Ovation Ultralow System V2 (TECAN, Redwood City, USA), following the manufacturer’s protocol. Ampure XP beads (Beckman Coulter) were used for final library purification. Library quality was analyzed on a 2200 TapeStation System with Agilent High Sensitivity DNA 1000 ScreenTape System (Agilent Technologies, Santa Clara, CA, USA). Resulting libraries were subjected to paired-end sequencing via Illumina HiSeq.

### 2.3 Metagenomic and metatranscriptomic bioinformatics

Metagenomic and metatranscriptomic data were annotated with the same pipelines. Briefly, adapter and primer sequences were filtered out from the Illumina paired reads, and then reads were quality trimmed to Phred33. rRNA reads were identified and removed with riboPicker [30]. We then assembled reads into transcript contigs using CLC Assembly Cell, and then we used FragGeneScan to predict open reading frames [ORFs; 31]. ORFs were functionally annotated using Hidden Markov models and blastp against PhyloDB [32]. Annotations which had low mapping coverage were filtered out (less than 50 reads total over all samples), as were proteins with no blastp hits and no known domains. For each ORF, we assigned a taxonomic affiliation based on Lineage Probability Index taxonomy [32, 33]. Taxa were assigned using two different reference databases: NCBI nt and PhyloDB [32]. Unless otherwise specified, we used taxonomic assignments from PhyloDB, because of the good representation of diverse marine microbial taxa.

ORFs were clustered by sequence similarity using Markov Clustering [MCL; 34]. Sequences were assigned MCL Clusters by first computing pairwise blastp for all sequences. The MCL algorithm was subsequently implemented using default parameters. MCL Clusters were then assigned consensus annotations based on KEGG, KO, KOG, KOG class, Pfam, TIGRfam, EC, GO, annotation enrichment [28, 32]. Proteins were assigned to coarse-grained protein pools (ribosomal and photosynthetic proteins) based on these annotations. For assignment, we used a greedy approach, such that a protein was assigned a coarse-grained pool if at least one of these annotation descriptions matched our search strings (we also manually examined the coarse grains for quality control). For photosynthetic proteins, we included light harvesting proteins, chlorophyll a-b binding proteins, photosystems, plastocyanin, and flavodoxin. For ribosomal proteins, we just included the term ‘ribosom*’ (where the * represents a wildcard character), and excluding proteins responsible for ribosomal synthesis.

### 2.4 Sample preparation and LC-MS/MS

We extracted proteins from the samples by first performing a buffer exchange from the sucrose-buffer to an SDS-based extraction buffer, after which proteins were extracted from each filter individually [as previously described 27]. After extraction and acetone-based precipitation, we prepared samples for liquid chromatography tandem mass spectrometry (LC-MS/MS). Precipitated protein was first resuspended in urea (100 *μ*L, 8 M), after which we measured the protein concentration in each sample (Pierce BCA Protein Assay Kit). We then reduced, alkylated, and enzymatically digested the proteins: first with 10 *μ*L of 0.5 M dithiothreitol for reduction (incubated at 60 C for 30 minutes), then with 20 *μ*L of 0.7 M iodoacetamide (in the dark for 30 minutes), diluted with ammonium bicarbonate (50 mM), and finally digested with trypsin (1:50 trypsin:sample protein). Samples were then acidified and desalted using C-18 columns [described in detail in 35].

To characterize each metaproteomic sample, we employed one-dimensional liquid chromatography coupled online to the mass spectrometer [VelosPRO Orbitrap, Thermo Scientific, San Jose, California, USA; detailed in 35]. For each injection, protein concentrations were equivalent across sample weeks, but different across filter sizes. We had higher amounts of protein on the largest filter size (3.0 um) and less on the smaller filters, so we performed three replicate injections per 3.0 *μ*m filter sample, and two replicate filter injections for 0.8 and 0.1 *μ*m filters. We used a non-linear LC gradient totalling 125 minutes. For separation, peptides eluted through a 75 *μ*m by 30 cm column (New Objective, Woburn, MA), which was self-packed with 4 *μ*m, 90 A, Proteo C18 material (Phenomenex, Torrance, CA), and the LC separation was conducted with a Dionex Ultimate 3000 UHPLC (Thermo Scientific, San Jose, CA).

### 2.5 LC-MS/MS bioinformatics – Database searching, configuration, and quantification

Metaproteomics requires a database of potential protein sequences to match observed mass spectra with known peptides. Because we had sample-specific metagenome and metatranscriptome sequencing for each metaproteomic sample, we assessed various database configurations, including those that we predict would be sub-optimal, to examine potential options for future metaproteomics researchers. We used five different configurations, described below. In each case, we appended a database of common contaminants (Global Proteome Machine Organization common Repository of Adventitious Proteins). We evaluated the performance of different database configurations based on the number of peptides identified (using a peptide false discovery rate of 1%).

In order to make these databases (Table 1), we performed three separate assemblies on 1) the metagenomic reads (from samples GOS-927, GOS-930, GOS-933 and GOS-935), 2) metatranscriptomic reads (from samples GOS-927, GOS-930, GOS-933 and GOS-935) and 3) metatranscriptomic reads from a concurrent metatranscriptomic experiment, started at the location where GOS-933 was taken [28]. Database configurations were created by subsetting from these assemblies. The first configuration was ‘one-sample database’, constructed to represent the scenario where only one sample was used for metagenomic and metatranscriptomic sequencing (we chose the first sampling week). Specifically, this was done by subsetting and including ORFs from the metagenomic and metatranscriptomic assemblies if reads from this time point were present in that sample [reads mapped as in 28], and then removing redundant protein sequences (P. Wilmarth, fasta utilities). The second configuration was the ‘sample-specific database’, where each metaproteomic sample had one corresponding database (prepared from both metagenome and metatranscriptome sequencing completed at the same sampling site), also done by subsetting ORFs from the metagenomic and metatranscriptomic assemblies as described above. The third configuration was pooling databases across size fractions – such that all metagenomic and metatranscriptomic sequences across the same filter sizes (e.g. 3.0 *μ*m) were combined. ORFs were subsetted from the metagenomic and metatranscriptomic assemblies as above. The fourth and fifth configurations are from the concurrent metatranscriptomic experiment[28]. The fourth configuration (‘metatranscriptome experiment (T0)’) was the metatranscriptome of the *in situ* microbial community (i.e. at the beginning of the experiment). This database was created by subsetting from the ‘metatranscriptome experiment (all)’ assembly. Finally, the fifth configuration was the metatranscriptome of all experimental treatments pooled together (two iron levels, three temperatures; ‘metatranscriptome experiment (all)’). The overlap between databases (potential tryptic peptides) in different samples is presented graphically in Supplementary Figs. S1, S2, S3.

**Table 1:** Characteristics of the five different database configurations we used for metaproteomic database searches. For the ‘One-Sample Database’, the second time point was used, and all samples were matched according to filter sizes. For the ‘Sample-Specific Databases’, each database was matched with the corresponding metaproteomic sample. For the ‘Pooled-Across-Sizes Databases’, databases were pooled across every time point and matched according to filter size. For these aforementioned databases, the metagenomic and metatranscriptomic proteincoding sequences were pooled. For the ‘Metatranscriptome Experiment (T = 0)’, only the first sampling point from the metatranscriptome experiment was included. For the ‘Metatranscriptome Experiment (all)’ configuration, all protein coding sequences were included from the treatment outcomes as well as the T = 0. *Averages are presented for Sample Specific Databases

| Database Configuration | Filter Size | Number of Protein Sequences in the Database* |
| --- | --- | --- |
| One-Sample Database | 0.1 | 664521 |
| One-Sample Database | 0.8 | 642132 |
| One-Sample Database | 3 | 334394 |
| Sample-Specific Databases | 0.1 | 713153* |
| Sample-Specific Databases | 0.8 | 633756* |
| Sample-Specific Databases | 3 | 440990* |
| Pooled-across-sizes Databases | 0.1 | 836620 |
| Pooled-across-sizes Databases | 0.8 | 855430 |
| Pooled-across-sizes Databases | 3 | 723057 |
| Metatranscriptome Experiment (T = 0) | 0.2 | 443681 |
| Metatranscriptome Experiment (all) | 0.2 | 2185747 |

After matching mass spectra with peptide sequences for each database configuration [MSGF+ with OpenMS, with a 1% False Discovery Rate at the peptide level; 36, 37], we used MS1 ion intensities to quantify peptides. Specifically, we used the FeatureFinderIdentification approach, which cross-maps identified peptides from one mass spectrometry experiment to unidentified features in another experiment – increasing the number of peptide quantifications [38]. This approach requires a set of experiments to be grouped together (i.e. which samples should use this cross-mapping?). We grouped samples based on their filter sizes (including those samples that are replicate injections). First, mass spectrometry runs within each group were aligned using MapAlignerIdentification [39], and then FeatureFinderIdentification was used for obtaining peptide quantities.

After peptides have been identified and quantified, we mapped them to proteins, which have corresponding functional annotations [KEGG, KO, KOGG, Pfams, Tigrfam; 28]. Functional annotations were used in three separate analyses. 1) Exploring the overall functional changes in microbial community metabolism, we mapped peptides to MCL clusters – groups of proteins with similar sequences. These clusters have consensus annotations based on the annotations of proteins found within the clusters [described in detail in 28]. For this section, we only used peptides that uniquely map to MCL clusters. 2) We restricted the second analysis to two protein groups: ribosomal and photosynthetic proteins. For this analysis, we mapped peptides to one of these protein groups if at least one annotation mapped to the protein group (via string matching with keywords). This approach is ‘greedy’ because does not exclude peptides if they also correspond with other functional groupings, but this is necessary because of the difficulties in comparing various annotation formats. 3) The last analysis for functional annotations was for targeted proteins, and we only mapped functions to peptides where the peptides uniquely identify a specific protein (e.g. plastocyanin).

Code for the database setup and configuration, database searching, and peptide quantification is open source (https://github.com/bertrand-lab/ross-sea-meta-omics).

### 2.6 LC-MS/MS bioinformatics – Normalization

Normalization is an important aspect of metaproteomics: it influences all inferred peptide abundances. Typically, the abundance of a peptide is normalized by the sum of all identified peptide abundances. We use the term normalization factor for the inferred sum of peptide abundances. Note that the apparent abundance of observed peptides is dependent on the database chosen. In theory, if fewer peptides are observed because of a poorly-matching database, this will decrease the normalization factor, and those peptides that are observed will appear to increase in abundance. It is not known how much this influences peptide quantification in metaproteomics.

For each database configuration, we separately calculated normalization factors. We then correlated the sum of observed peptide abundances with each other. To get a database-independent normalization factor, we used the sum of total ion current for each mass spectrometry experiment [using pyopenms; 40], and also examined the correlation with database-dependent normalization factors. If normalization factors are highly correlated with each other, that would indicate database choice does not impact peptide quantification.

### 2.7 Defining proteomic mass fraction

Protein abundance can be calculated in two ways: 1) the number of copies of a protein (independent of a proteins’ mass), or 2) the total mass of the protein copies (the sum of peptides). We refer to the latter as a proteomic mass fraction. For example, to calculate a diatom-specific, ribosomal mass fraction, we sum all peptide abundances that are diatom- and ribosome-specific, and divide by the sum of peptide abundances that are diatom-specific. Note that this is slightly different to other methods, like the Normalized Spectral Abundance Factor, which normalizes for total protein mass [via protein length; 41].

### 2.8 Combining estimates across filter sizes

Organisms should separate according to their sizes when using sequential filtration with decreasing filter pore sizes. In practise, however, organisms can break because of pressure during filtration, and protein is typically present for large phytoplankton on the smallest filter size and vice-versa. We used a simple method for combining observations across filter sizes, weighted by the number of observations per filter. We begin with the abundance of a given peptide observed, which was only considered if it was observed across all injections, and was then taken as the average abundance across injections. We calculated the sum of observed peptide intensities (i.e. the normalization factor), and divided all peptide abundances by this normalization factor. If we are estimating the ribosomal mass fraction of the diatom proteome, we first normalize the diatom-specific peptide intensities as a proportion of diatom biomass (i.e. divide all diatom-specific peptides by the sum of all diatom-specific peptides). We then summed all diatom-normalized peptides intensities that are unique to both diatoms and ribosomal proteins, which would give us the ribosomal proportion of the diatom proteome. Yet, we typically would obtain multiple estimates of, for example, ribosomal mass fraction of diatoms, on different filters. We combined the three values by multiplying each by a coefficient that represents a weight for each observation (specific to a filter size). These coefficients sum to one, and are calculated by summing the total number of peptides observed at a time point for a filter, and dividing by the total number of peptides observed across filters (but within each time point). For example, if we observed 100 peptides that are diatom- and ribosome-specific, and 90 of these peptides were on the 3.0 *μ*m filter and only 10 were on the 0.8 um filter, we would multiply the 3.0 *μ*m filter estimate by 0.9 and the 0.8 *μ*m filter by 0.1. This method uses all available information about proteome composition across different filter sizes [similar to 42].

When we estimate the proteomic mass fraction of a given protein pool, we do not need to adjust for the total protein on each filter. This is because this measurement is independent of total protein. However, for merging estimates of total relative abundance of different organisms across filters, we needed to additionally weight the abundance estimate by the amount of protein on each filter. Therefore, in addition to the weighting scheme described above, we multiplied taxon abundance estimates by the total protein on each filter divided by the total protein across filters on a given day.

### 2.9 LC-MS/MS simulation

We used simulations of metaproteomes and LC-MS/MS to 1) quantify biases associated with inferring coarsegrained proteomes from metaproteomes, and 2) to mitigate these biases in our inferences. Specifically, we asked the question: how does sequence diversity impact quantification of coarse-grained proteomes from metaproteomes? Consider a three organism microbial community. If two organisms are extremely similar, there will be very few peptides that can uniquely map to those organisms, resulting in underestimated abundance. A similar outcome is anticipated with differences in sequence diversity across protein groups, such that highly conserved protein groups will be underestimated.

Our mass spectrometry simulations offer a unique perspective on this issue: we know the ‘true’ metaproteome, and we can compare this with an ‘inferred’ metaproteome. We simulated variable numbers of taxonomic groups, each with different protein pools of variable sequence diversity. From this simulated metaproteome, we then simulated LC-MS/MS-like sampling of peptides. Complete details of the mass spectrometry simulation are available in [43] and the supplementary materials. The only difference between this model and that presented in [43] is here we include dynamic exclusion.

### 2.10 Cofragmentation bias scores for peptides

We recently developed a computational model (‘cobia’) that predicts a peptides’ risk for interference by sample complexity [more specifically, by cofragmentation of multiple peptides; 43]. This study showed that coarsegrained taxonomic and functional groupings are more robust to bias, and that this model can also be used to estimate bias. We ran cobia with the sample-specific databases, which produces a ‘cofragmentation score’ – a measure of risk of being subject to cofragmentation bias. Specifically, the retention time prediction method used was RTPredict [44] with an ‘OLIGO’ kernel for the support vector machine. The parameters for the model were: 0.008333 (maximum injection time); 3 (precursor selection window); 1.44 (ion peak width); and 5 (degree of sparse sampling). Code for running this analysis, as well as the corresponding input parameter file, is found at https://github.com/bertrand-lab/ross-sea-meta-omics.

## 3 Results and Discussion

### 3.1 Assessing database configurations and consequent quantification

The sequence databases from the metatranscriptome experiment conducted on our third sampling week (January 15, 2015) outperformed sample-specific databases and other configurations (in terms of number of peptide spectrum matches, Supplementary Fig. S4; Table 1). Specifically, we identified 14 455 unique peptides using the ‘metatranscriptome experiment T0’ database, while 8022 unique peptides were identified with the ‘sample-specific database’ (Supplementary Fig. S4). 5127 peptides were identified as a core set, regardless of the database chosen (Supplementary Fig. S4). The database pooled across time points identified more peptides than the ‘sample-specific database’, similar to previous work [45]. The metatranscriptomic experiment (both ‘metatranscriptomic experiment (T0)’ and ‘metatranscriptomic experiment (all)’) were more valuable in identifying larger, primarily eukaryotic organisms (Supplementary Fig. S4, S5, S6, S7). Overall, the two metatranscriptomic experiment databases performed similarly in terms of number of identified peptides. All subsequent analyses use the identified peptides from the ‘metatranscriptome experiment (all)’ database. Importantly, a difference between the metatranscriptomes of sample-specific filters and the metatranscriptomic experiment databases was sequencing depth (Supplementary Table 1). This difference likely influenced the metatranscriptomic read assembly, improving the assembly of eukaryotic-protein sequences and therefore creating a better database (i.e. in terms of peptides identified). Note that databases were constructed *after* assembly, and then subsetted to create individual databases (Methods). Overall, deep metatranscriptomic sequencing appears to be a promising avenue for metaproteomics with tailored databases (Supplementary Table 1).

Database choice influenced peptide quantification due to normalization. We quantified this by correlating sample-specific normalization factors with each other and with the total ion current (i.e. a database-independent normalization, Supplementary Figs. S8, S9, S10). Examining the correlation between the best- and worst-performing databases, there was a range of R^2^ values, from 83–99% (Fig. Supplementary Figs. S8, S9, S10). If we consider a peptide observed in a mass spectrometry experiment with an intensity value of 100, we expect variation in the inferred value to range from 92 to 108, reflecting variation in the normalization factor of 16%. This has significant consequences for comparative metaproteomics: consider two samples, one with a perfectly matched database and a second that uses the same database, but is poorly matched. Using standard methods, the peptides identified in the second sample will appear to increase in abundance, even if the abundance is constant. Note that our worst performing database was still well-matched to the community, so for researchers studying very distinct communities it is vital to address this issue by using database-independent normalization, or ensuring bias across samples is minimal. We provided methods for doing both.

One simple alternative is to use a database-independent metric of total peptide abundance: total MS1 ion current (TIC). We found that TIC is well correlated with the total peptide abundance inferred from the best-performing database (Supplementary Fig. S8, S9, S10). This result has two consequences: 1) it suggested that TIC may be a viable alternative for normalization in comparative metaproteomics. 2) it validated the use of our best-performing database, as we identified most of the abundant peptides in our sample. Given that these two approaches were highly correlated, we used the ‘metatranscriptome experiment (all)’ database for all subsequent analyses.

### 3.2 Characterizing eukaryotic-containing metaproteomes and quantifying unknown protein groups

Taxonomic abundance shifted through the season at the Antarctic sea ice edge (Fig. 1a). The microbial community was dominated by *Phaeocystis antarctica* (Haptophyta) early in the season, with diatoms increasing in relative abundance later on (predominantly *Fragilariopsis* sp. and *Pseudonitzschia* sp.). The phytoplankton bloom progression and high dinoflagellate biomass contribution were both consistent with previous observations in the Ross Sea [46, 47].

**Figure 1:**
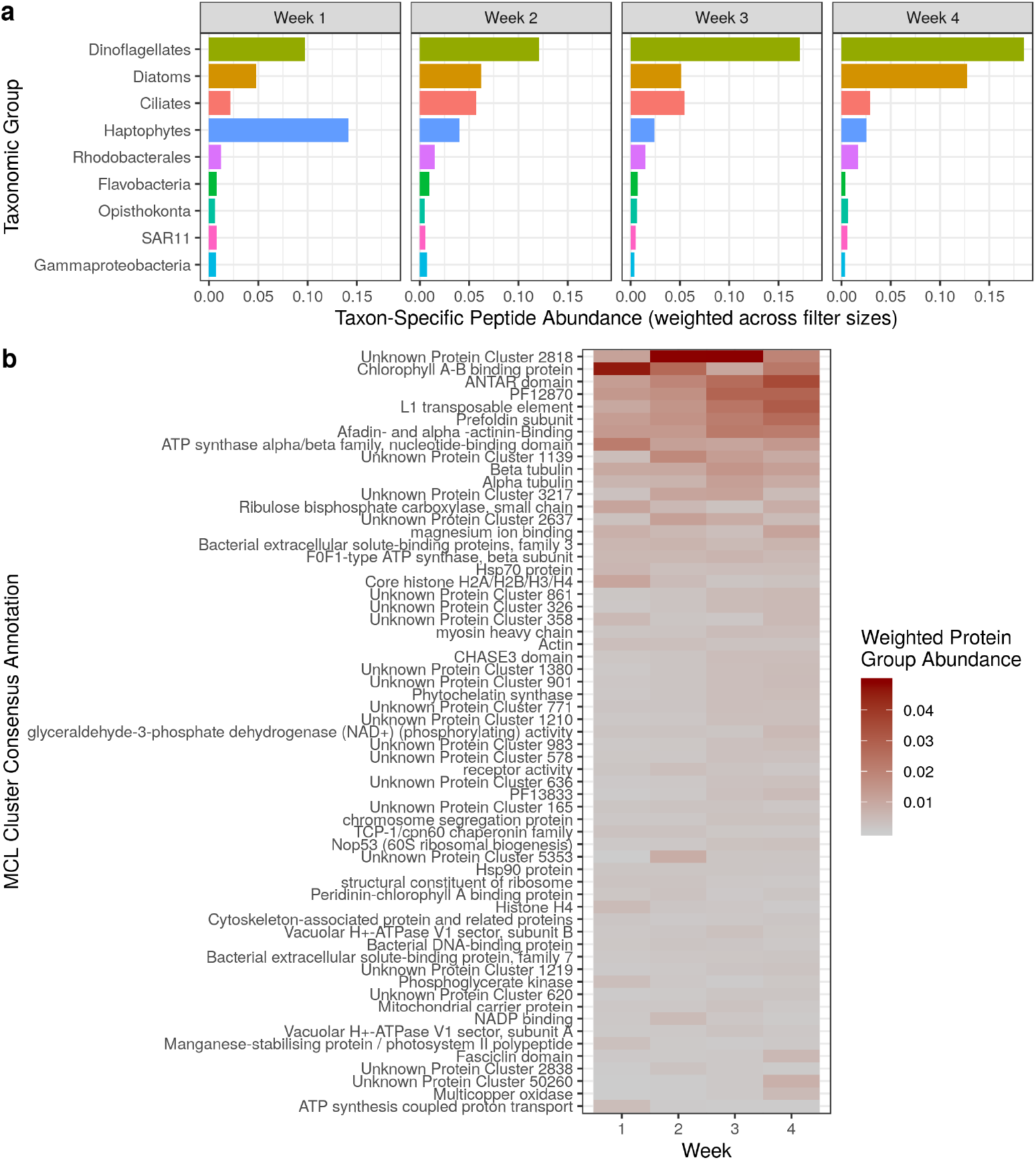
**a**, Measurements of relative change in protein biomass identified a taxonomic shift at the Antarctic sea ice edge. Protein biomass is calculated as the sum of taxon-specific peptide intensities, weighted by the protein mass per filter for each sampling time (Methods). **b**, relative change in protein functional clusters shows that unknown protein clusters contribute greatly to in situ protein biomass, and also identifies a functional shift across weeks.

We identified functional shifts in protein abundance by mapping peptides to de novo protein clusters (independent of taxonomic assignments) – including protein clusters with no known function. Earlier in the season there was a high relative abundance of Chlorophyll A-B binding proteins and ATP synthase alpha/beta family proteins (Fig. 1b), which is anticipated because of the higher levels of dissolved iron [27]. Demonstrating the importance of de novo protein group assignment, the most abundant protein group in our entire dataset had no functional annotations (Fig. 1b, Unknown Protein Cluster 2818, mostly belonging to Ciliates). Further examination of a protein sequence within this cluster found no functionally similar proteins within the NCBI Non-redundant database. We suggest that these unknown, highly abundant proteins should be targets for functional characterization.

### 3.3 Eukaryotic and prokaryotic taxa have taxon-specific proteomic allocation strategies

We quantified two simple proteomic ‘traits’ of microbes: the ribosomal protein mass fraction and the photosynthetic protein mass fraction. Eukaryotic taxa formed unique clusters based on these two traits, with more variation across taxa than across time points (Fig. 2a). For example, haptophytes however had relatively high proportions of both ribosomal and photosynthetic protein fractions. Examining the five most abundant prokaryotic taxa, we also observed distinct proteomic compositions, with gammaproteobacteria exhibiting the highest ribosomal protein mass fraction (Fig. 2).

**Figure 2:**
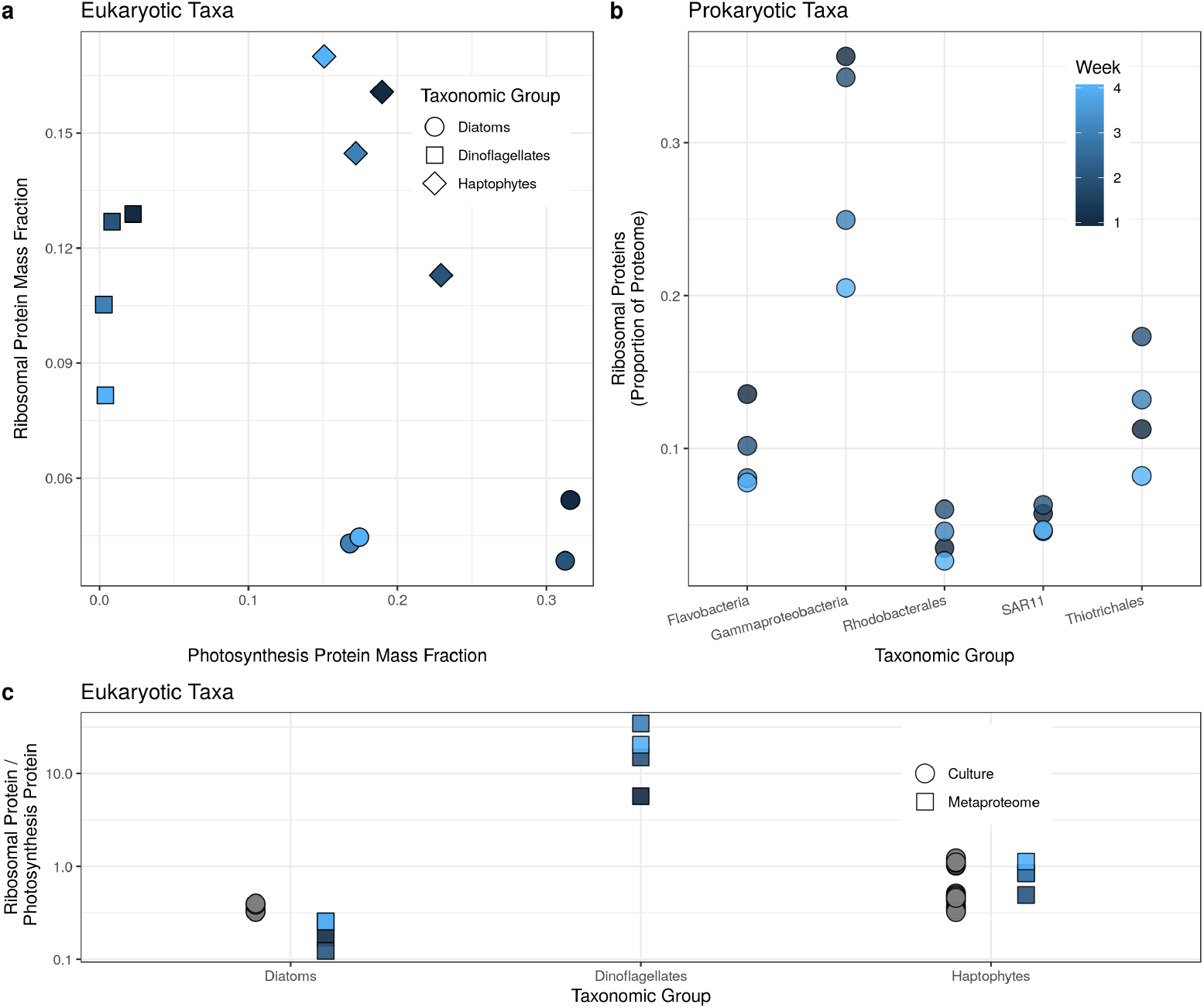
**a**, photosynthetic protein mass fraction and ribosomal protein mass fraction (normalized by total amount of a given taxon at a each time point) identifies clear taxonomic subgroupings. **b**, examining prokaryotic taxa only shows variation in ribosomal mass fraction across groups. **c**, Ratios of ribosomal protein mass fraction to photosynthetic protein mass fraction derived from metaproteomic observations and compared with phytoplankton proteomes observed in culture [*Phaeocystis antarctica*, *Thalassiosira pseudonana* 27, 57].

Before examining the underpinnings of these proteomic traits, we first scrutinized these inferences using mass spectrometry simulations and additional data sources. Our analysis was limited to coarse-grained protein functions and taxa, which is robust to bias arising from variable sample complexity [43]. Our mass spectrometry simulations suggested that low sequence diversity in taxa or protein groups can lead to underestimation (Supplementary Fig. S12), but this bias is mitigated by examining abundant proteins or taxa. 100 to 400 taxon-specific peptides were identified for each protein group, and our simulations indicated that this is sufficient to avoid significant bias. We therefore restricted our analyses to taxa and protein groups that are relatively abundant. Note that for dinoflagellates, we observed relatively few peptides in the photosynthetic proteomic mass fraction, so our observations are likely underestimating the true value (Supplementary Discussion, Supplementary Fig. S12). Despite this underestimation, the true value is probably quite low (discussed below).

We provided two additional estimates of ribosomal and photosynthetic protein mass fraction from cultured phytoplankton (Fig. 2c). Metaproteomics can underestimate taxon-specific protein mass when taxonomically uninformative peptides are not used. For example, we might identify a highly conserved peptide produced by a diatom, but are unable to map it to diatoms because it also corresponds to other taxa, and this peptide would be excluded from the quantification of diatoms. Therefore, we compared the ratios of ribosomal to photosynthetic protein mass fraction from the metaproteomic observations to cultured diatoms and haptophytes. Ratios were similar in cultures compared to populations sampled in situ (Fig. 2c), despite such culturing experiments occurring under different environmental conditions. Trends observed in the proportion of transcripts mapped to ribosomal proteins in different groups of prokaryotes also mirrored our estimates of ribosomal protein mass fraction [high for gammaproteobacteria and low for SAR11; 21].

We examined coarse-grained taxonomic groups. It is possible that within these coarse groupings, different taxa included in these groupings employ different allocation strategies. We therefore sought to determine whether taxonomic sub-groupings displayed similar expression patterns. This issue is challenging to assess, because as subgroupings are further examined, there is increased susceptibility to several biases (as outlined above). We therefore examined one subgrouping, diatoms, that contained two dominant species: *Fragilariopsis sp*. and *Pseudonitzschia sp*. The taxonomic assignments for these two diatoms were from the NCBI nt database. We observed similar proteome estimates for both ribosomal and photosynthetic proteins amongst both these subgroups of diatoms (Supplementary Fig. S13), suggesting they are functionally similar based on these proteomic traits. However, we cannot exclude the possibility that for other taxonomic groups the trends observed are due to a diversity of underlying microbial strategies. Yet at this coarse taxonomic level, we concluded that different microbial taxa exhibited distinct coarse-grained proteomes.

We now turn to the ecological relevance of these protein expression patterns. Protein synthesis is the primary energy sink in cells [25], and photosynthesis or respiration is the primary energy source in cells. Why do dinoflagellates have relatively low photosynthetic protein mass fractions? This taxonomic group is typically mixotrophic or heterotrophic, which would require larger investment in respiratory proteins for energy production. Haptophytes and diatoms had similar amounts of photosynthetic proteins, but very different amounts of ribosomal proteins (Fig. 2a), so there was no direct trade-off between producing ribosomal versus photosynthetic machinery [i.e. they do not form Pareto front; 48, 49]. Gammaproteobacteria had the highest ribosomal mass fraction within the observed prokaryotic taxa, and haptophytes had higher ribosomal mass fractions compared to diatoms.

What are the ecological implications of having more ribosomes? If we assume constant translation rate per translational apparatus [but see 50], taxa then had different total protein synthesis output. Growth rate is directly related to total protein synthesis output, because protein comprises a large portion of cell mass. To have a faster growth rate, microbes’ need to increase protein synthesis [see 12, for derivation and assumptions]. We hypothesize that high total protein synthesis output (via high ribosomes) is more advantageous under high nutrient regimes, as it would allow an elevated growth rate. Indeed, haptophytes and gammaproteobacteria were more abundant earlier in the season [which had higher concentrations of dissolved Fe and Mn; 27]. Another interpretation is that these early-abundant taxa are better suited to a dynamic environment. Perhaps these early-abundant taxa (gammaproteobacteria, haptophytes) increased investment in ribosomes as a form of bet-hedging, which enables a faster growth rate in a dynamic environment [51].

### 3.4 Environment-independent proteomic fraction varies across taxa

What is the cost of responding quickly to a dynamic environment? We hypothesized that there is a regulatory cost for producing proteins that are optimal for a set of environmental conditions. Constitutive protein production does not incur this regulatory cost at the risk of being mismatched to environmental conditions. If the proteome is mostly constant across conditions, this indicates a low regulatory cost, and vice versa. We propose a proteomic trait that reflects regulatory cost: the proteomic fraction that is environment-independent. This proteomic trait is quantifiable using metaproteomics, and due to the dynamic nature of the ocean, is likely an important selective force for marine microbes.

We classified peptides that are ‘constant’ across different environmental conditions, and then summed their average intensities to get an environment-independent peptide mass fraction (Fig. 3b and c). Note that 1) peptide intensities were first normalized by total taxon-specific peptide intensity (they therefore sum to one for each taxon), and 2) estimates of environment-independent peptide mass fraction were combined across filter sizes (Methods). Using proteomic data from replicate cultures under identical conditions, we chose a cut-off point distinguishing environment-dependent versus-independent peptides [represented with vertical lines, Fig. 3a; Supplementary Fig. S15; 52]. We then can determine the proportion of the proteome that is environment-independent and – dependent (using the mean abundance value per peptide). There are potential biases in this novel method. We address the impact of these biases using published data and by making comparisons with other estimates of regulatory costs across taxa from previously published work (see Supplementary Discussion).

**Figure 3:**
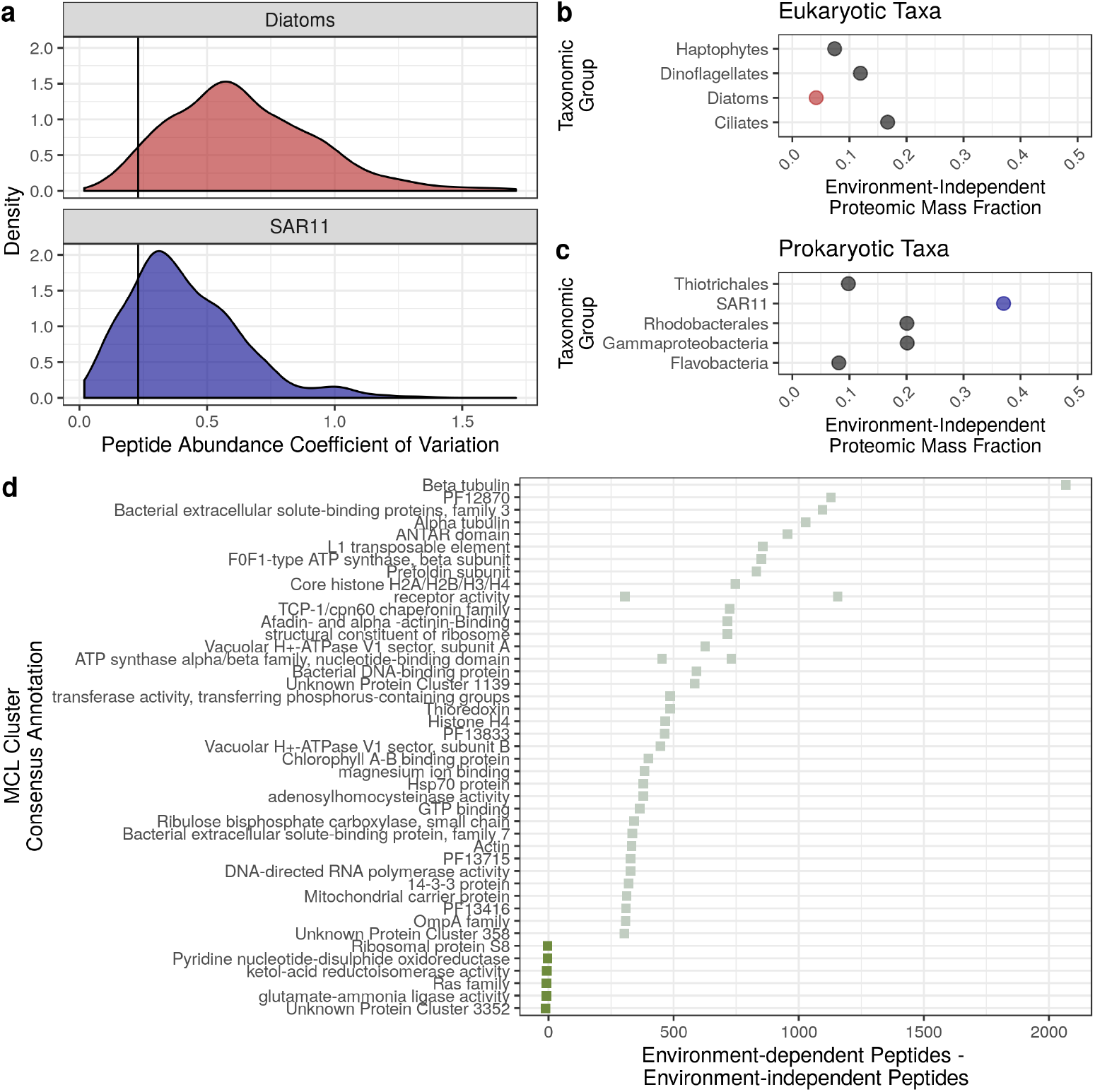
**a**, the distribution of peptide-specific coefficients of variation can be used to identify if a peptide is significantly changing across environmental conditions. Peptide abundance is first divided by the total taxon-specific peptide intensity. Diatoms and SAR11 represent two extremes within this dataset – diatoms have a highly variable proteome while SAR11 has a relatively constant protein expression. The cutoff point was chosen using replicate cultures of a diatom proteome [vertical line 52]. **b, c**, after classifying peptides by their coefficients of variation, we categorized peptides as independent of their environment and those that are not. Points represent the sum of peptide intensities that are environment independent across eukaryotic and prokaryotic taxa. **d**, a comparison of protein functional clusters that are mostly either environment-dependent or environment-independent. Positive values indicate that peptides observed within this protein cluster were mostly dependent on their environment (high coefficient of variation), while negative values indicating most peptides within this protein cluster were identified as environment-independent.

SAR11 had the highest environment-independent peptide mass fraction across all eukaryotic and prokaryotic taxa we examined (Fig 3a, 3b), consistent with previous work suggesting SAR11 has reduced regulatory investment [53]. Within eukaryotes, dinoflagellates exhibited the highest environment-independent peptide mass fraction, and dinoflagellates in other oceanic regions also exhibited lower regulatory cost [20, 22].

Diatoms had a lower environment-independent proteomic fraction compared with haptophytes, suggesting they have higher regulatory costs. Recall the previous result that diatoms had a lower proportion of ribosomes compared with haptophytes (but similar proportions of photosynthetic proteins; Fig. 2a). We speculate that two proteomic traits comprise a trade-off for these two taxa: higher total protein synthesis via more ribosomes (i.e. leading to fast growth under high nutrient conditions), but at a cost of being less able to dynamically regulate their proteomes. This suggests that in a high nutrient environment (that is also dynamic), dynamically responding to the environment is not the optimal strategy. Instead, a better strategy is constitutively expressing proteins that are favourable for rapid growth (e.g. high ribosomal production in haptophytes), which may help explain the progression from haptophyte-to diatom-dominated communities in the Ross Sea bloom.

Are some protein functions more often categorized as environment-independent or environment-dependent? Highlighting some examples, the actin protein cluster was often classified as environment-dependent (Fig. 3c). Actin is involved in endocytosis, and inorganic Fe uptake occurs via an endocytotic mechanism [with phytotransferrin; 54]. Perhaps variable expression of actin is related to the amount of bioavailable Fe, and previously published proteomic experiments also showed that actin was differentially expressed due to Fe [55, 56]. ATP synthase-peptides and chlorophyll A-B binding protein-peptides were also mostly classified as environment-dependent, likely reflecting higher primary production earlier in the season (Fig. 3c). In contrast, the ketol-acid reductoisomerase protein cluster (involved in branched-chain amino acid synthesis) was mostly classified as environment-independent. It is unclear what the mechanistic basis for constitutive expression of this protein might be, but several proteomic studies of diatoms also suggest similar expression across conditions [55, 56, 57]. Using this extensible approach to identify constitutively expressed proteins across a wide array of taxa would shed light on these mechanisms. With vastly more metaproteomic data being generated [e.g. 58], identifying constitutively expressed proteins across diverse taxa would help answer the question: what are the features of constitutively expressed proteins?

### 3.5 Coarse-grained proteomes for measuring nutrient stress

Proteomics is also used in marine microbiology to assess stress corresponding to a deficient nutrient [e.g. 4]. For example, expression of the protein plastocyanin may reflect Fe deficiency, because plastocyanin does not contain Fe and performs a similar function as the Fe-containing protein cytochrome *c* [59]. Biomarkers of physiological stress are increasingly nuanced [e.g. 27], sometimes taxon specific, and can require targeted mass spectrometry approaches. Coarse-grained approaches may be a complementary method for assessing stress or nutrient deficiency. We compared using coarse-grained proteomes with single-protein biomarkers. We first solely examined the photosynthetic protein mass fraction compared to the mass fraction of peptides assigned to the plastocyanin, for diatoms and haptophytes (Fig. 4a-b). This approach is biased by variable complexity across samples [43], but we predicted the degree of bias with a quantitative metric (the ‘cofragmentation score’). This score reflects the expected number of peptides with similar m/z and retention times. Overall, there were relatively few potential cofragmenting peptides (≈3), indicating low bias (peptides with high bias can have upwards to 300 cofragmenting peptides, for example [43]. We observed a negative relationship between the photosynthetic protein mass fraction and the plastocyanin mass fraction (note that these two variables are not independent, as plastocyanin is considered as part of the photosynthetic mass fraction). We also examined Phaeocystis antarctica-specific peptides measured with targeted mass spectrometry, and identified a negative correlation between the abundance values of plastocyanin and the coarse-grained estimates of photosynthetic proteins (Fig. 4c). We conducted this analysis as a proof-of-concept for using coarse-grained proteomes to assess nutrient deficiency, as coarse-grained proteomes are amenable for untargeted metaproteomic analyses. More analyses are required to assess the robustness of this relationship, and also to assess if coarse-grained proteomic signatures are nutrient specific.

**Figure 4:**
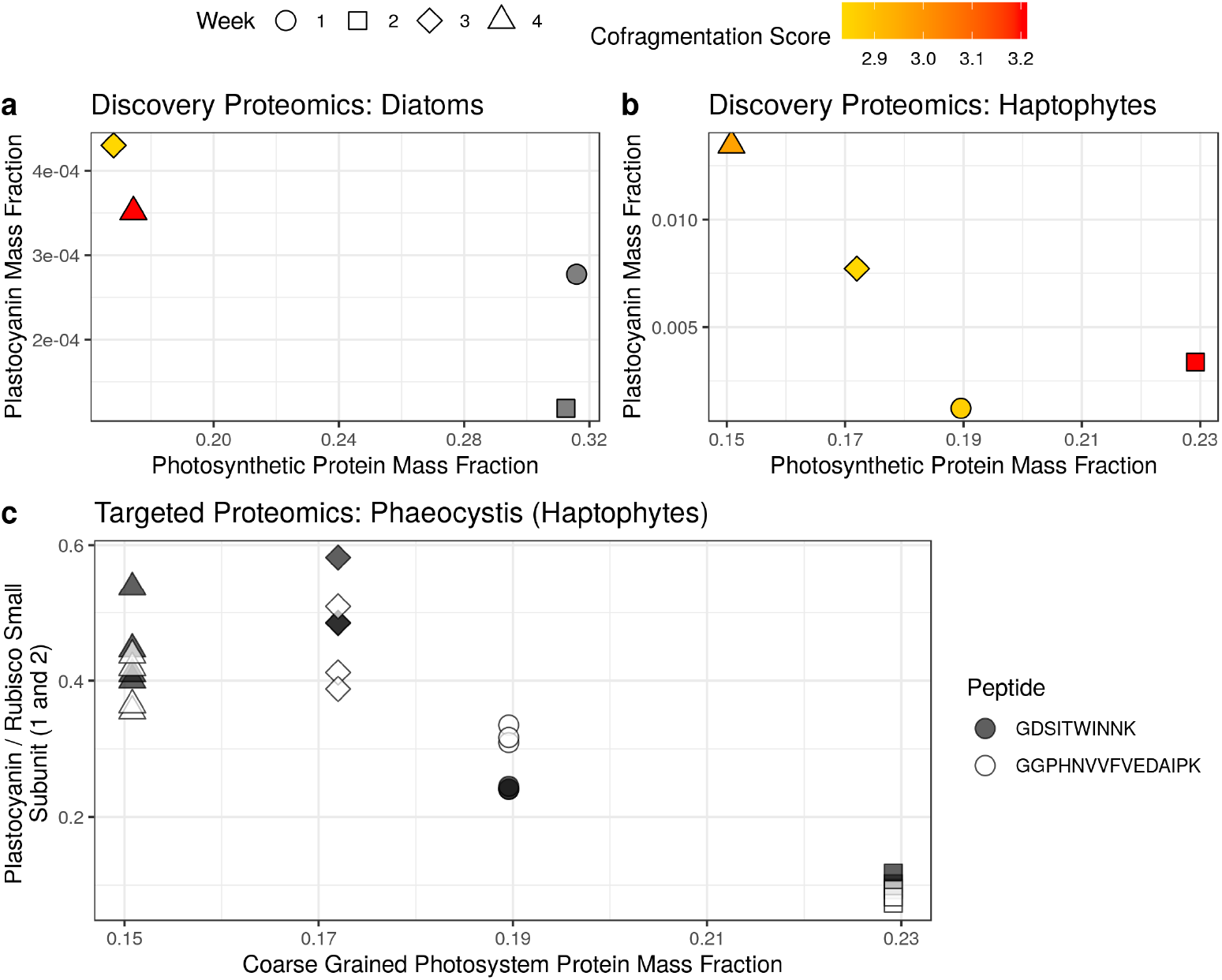
a-b, comparison of the single-protein biomarker (using untargeted, discovery proteomics) plastocyanin with the photosynthetic protein mass fraction for diatoms and haptophytes. c, comparison of the single-protein biomarker (using targeted proteomics) plastocyanin with the photosynthetic protein mass fraction for haptophytes. Two peptides for plastocyanin are shown. Plastocyanin abundance is normalized to Rubisco small subunit 1 and 2, each which had a taxon-specific peptide (AKPNFYVK and QIQYALNK respectively).

## 4 Conclusion

We conclude that different microbial taxa have distinct coarse-grained proteomic composition, and this composition is more similar across taxa than across environmental conditions. The stoichiometry of proteins within pathways is conserved [9] – but our results show that this is not the case across pathways. Variation in pathway-to-pathway stoichiometry may indeed underpin ecological strategies, in addition to differing gene repertoires. Connecting in situ proteomes to ecological strategies will delineate proteomic ‘traits’, which can then be adopted into a trait-based approach for modelling microbial communities. Genomic trait-based approaches have successfully explained large-scale biogeochemical processes [10, 11], but they first had to identify genes that are metabolically important. Therefore, identifying and quantifying proteomic trait variation across taxa will connect protein production to ecological strategies, and ultimately enable modelling of microbial communities by representing proteomic traits and trade-offs in large scale models [e.g. as in 60].

## 5 Data Availability

The metagenomics and metatranscriptomics data reported here have been deposited in the NCBI sequence read archive (BioProject accession no. PRJNA074702; BioSample accession nos. SAMN18057468-SAMN18057479 (metagenomics) and BioSample accession nos. SAMN18057480-SAMN18057497 (metatranscriptomics). Raw mass spectrometry data are deposited in the PRIDE database with project accession PXD022995 (accessible using reviewer username and password njAogAv6). All other data products (the ‘cobia’ analysis output, formatted databases, peptide abundances for each database search, targeted proteomics data, culture proteomics data, metaproteomic simulation output) are available in Dryad at doi:10.5061/dryad.vt4b8gtrz (access prior to publication: https://datadryad.org/stash/share/yQRc5qjdnE6ix-he5Bkuo0c42r4G4bSY-JzeSVIYYsM).

## 6 Acknowledgements

We thank David Hutchins, Jeff Hoffman, Rachel Sipler, Jenna Spackeen, and Deborah Bronk, as well as Antarctic Support Contractors and the staff at McMurdo Station for support in the field. We are grateful to Alejandro Cohen from the Dalhousie Biological Mass Spectrometry Core Facility for assistance with mass spectrometry data acquisition, and to Noor Youssef for discussion. J.S.P.M. acknowledges support from the NSERC CREATE Transatlantic Ocean System Science and Technology Program and the Killam Scholarship. This project was financially supported by NSERC Discovery Grant RGPIN-2015-05009 to E.M.B., Simons Foundation Grant 504183 to E.M.B., an NSERC CGS Postgraduate scholarship to J.S.P.M., NSF-ANT-1043671, NSF-OCE-1756884, and Gordon and Betty Moore Foundation Grant GBMF3828 to A.E.A.

## 10 Supplementary Methods

We simulated metaproteomes *in silico* to examine biases arising from inferring taxon-specific proteomes. The primary challenge of inferring taxon-specific coarse grained proteomes is that not all coarse grained pools (groups of proteins performing some functional role) are equally identifiable. This also extends to taxa – some taxa are more closely related, and therefore have fewer unique peptides. For example, some coarse grained pools are easily mapped to a given taxon while others have very few taxon-specific peptides.

To address these expected biases, we created *in silico* metaproteomic datasets (generative model), and sampled the data similar to how a mass spectrometer would (sampling model). We then compared the sampled data to the known dataset, and evaluated which conditions biases would arise.

### 10.1 Generative Model

We generate *p* unique peptides, assigned to *k* coarse grained pools, belonging to an organism *j*. We simulate peptides rather than proteins, as peptides are injected into a mass spectrometer with bottom-up mass spectrometry. To simulate different levels of sequence diversity present across protein pools, we generate k sequence ‘banks’ of different sizes. Peptide sequences banks are created by randomly sampling from all amino acids, generating a sequence ranging in length from 5–15 amino acids per peptide. An organism-specific peptide profile is created, which randomly samples from each ‘sequence bank’. So a smaller ‘sequence bank’ would represent a coarse grained protein pool with low diversity, and vice-versa.

We then assign abundances to each peptide. Peptide abundance is generated using a random sample from a gamma distribution with the shape parameter of 0.15 and the scale parameter of 10. We chose this distribution as it is similar to the distribution of peptides observed in single-organism proteomics (specifically it has overdispersion, non-zero values only, and is continuous). We then multiply each peptide abundance by a taxonomic abundance unique to each taxon *j*, and by the abundance within a given coarse grained pool *k*. Both the taxonomic abundance and the coarse-grained pool abundance values are similarly drawn from a gamma distribution, except with a shape value of 1. To arrive at a final abundance of a peptide we multiply the organism abundance (e.g. 100) by the abundance of given coarse-grained pool (e.g. 5). Lastly, we generate a value for all peptides from within this organism and coarse-grained pool (e.g. 2), and multiply these three values. In this case, the intensity of the peptide would be 1000. Once the ‘true’ dataset is generated, we then filter this dataset to create an ‘observed’ dataset, because peptides that are the same from the ‘true’ dataset should be summed. From this observed dataset, we calculate peptide mass and assume a peptide charge state of 2.

### 10.2 Sampling Model

Mass spectrometers sample and fragment peptides for identification, and there is some stochasticity in this sampling process, particularly when using data-dependent acquisition (DDA). Using DDA, peptides are sampled according to their intensity. We subsample our ‘observed dataset’ using a simplistic model of a mass spectrometer. Our model assumes a constant ion peak width, and randomly assigns elution times to peptides from a uniform distribution. A similar version of this model has been extensively validated ([43]), but the key difference here is including dynamic exclusion and top-N sampling.

We describe sampling model algorithmically below (Algorithm 1). We begin by sorting and then binning elution times for all peptides (steps 1–2). We then loop through every *n*th elution time bin, where *n* represents the number of ions selected for Top *n* DDA (step 4). So with more ions selected for ‘fragmentation’ the mass spectrometer would have less time to scan intact peptides, as is true for instruments that move between scanning MS1 in an Orbitrap and fragmenting peptides in a linear ion trap. Then, if a peptide is on the dynamic exclusion list and it has been on the list for longer than the dynamic exclusion time, it is removed from the dynamic exclusion list (step 4–5). All of the *m*/*z* windows belonging to a peptide on the dynamic exclusion list are then blocked for sampling (steps 6–7). Of the remaining peptides, we select the top *n* in terms of abundance, and assume that these peptides are identified (step 8). The final step is adding the identified peptides to the dynamic exclusion list (step 9), which prevents those *m*/*z* regions from being subsequently sampled for a short period of time (the dynamic exclusion time).

**Algorithm 1:**
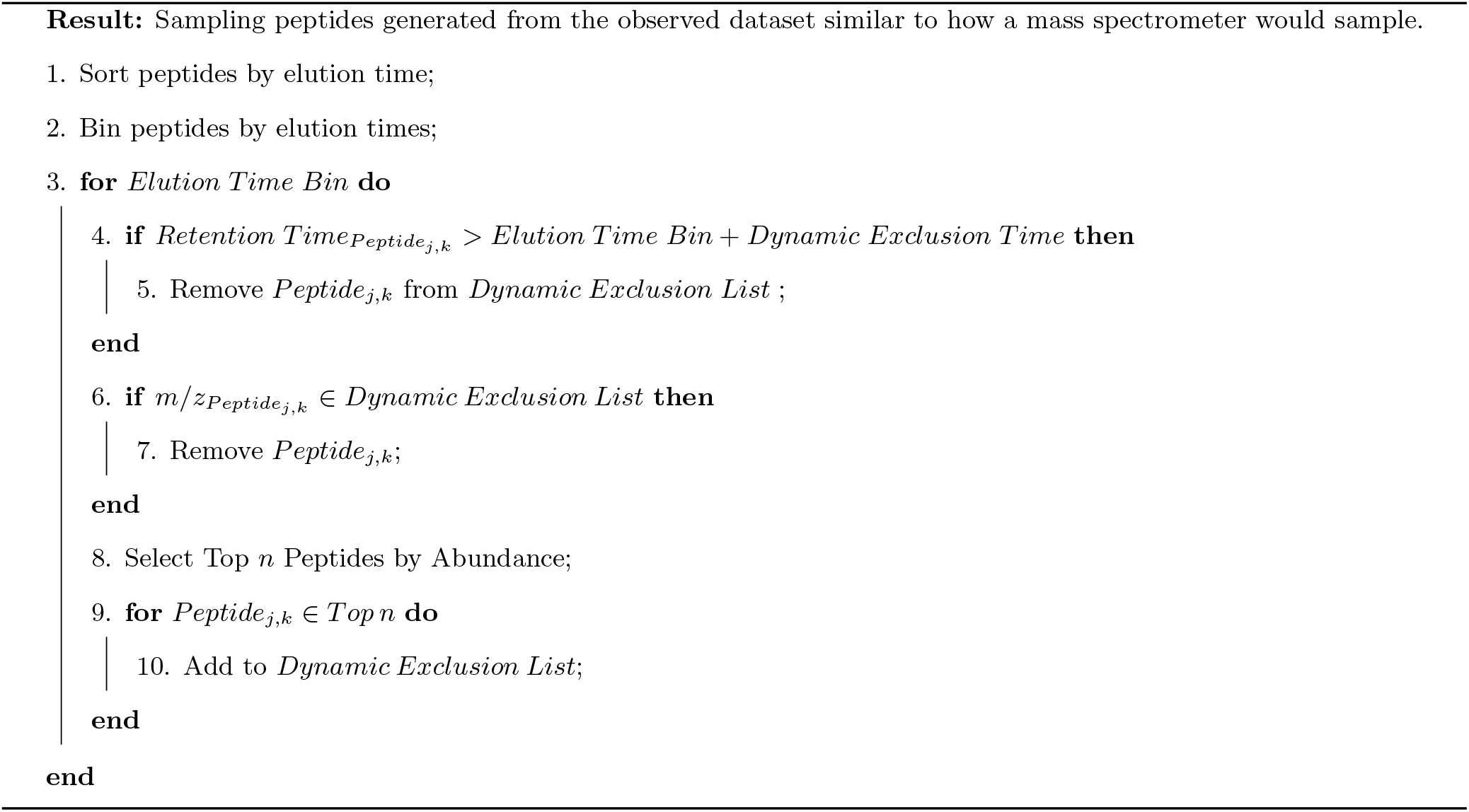
Mass spectrometry sampling model.

### 10.3 Model Parameters

We generated 15 datasets with the following characteristics. Each dataset contained 30 distinct taxa with four coarse-grained protein groups of varying diversity. As above, diversity is modeled using varying sizes of sequence ‘banks’ (we used sizes of 50000, 100000, 250000, and 500000). From each protein group (represented by these different sequence banks of peptides), each organism has 2000 peptides, which are randomly drawn from these sequence banks.

Retention times are assigned from a uniform distribution ranging from 0–90 minutes. The maximum injection time, which is used as the width of the elution time bin, is 500ms (or 0.00833 minutes), following from [43]. We assign a constant ion peak width of 0.5 minutes, independent of ion intensity. We use a Top *n* of 12 ions. Our precursor selection window is set to 3*m*/*z* and our dynamic exclusion time span is set to 0.5 minutes.

## 11 Supplementary Discussion

### 11.1 Underestimation of coarse-grained protein groups

Our simulations showed that abundant protein groups have good estimates (close to the 1:1 line, Supplementary Fig. S12), while low abundance protein groups tend to be underestimated. This is because of the data-dependent acquisition sampling method that mass spectrometers use. Data-dependent acquisition specifically targets the highly abundant peptides, so lower abundance groups tend to get sampled less. The method we (and others) typically use is to sum the peptide intensities to obtain an abundance estimate. With fewer peptides quantified, the sum will be lower (Supplementary Fig. S12). Note that sequence diversity can also influence these estimates (represented with blue colour gradient, Supplementary Fig. S12), but only until there is extremely low diversity (darkest colour), corresponding with only a few peptides identified and mapped to a taxon.

### 11.2 Sequence Diversity

Our conclusions about the ribosomal and photosynthetic proteomic mass fractions, as well as the environment-independent proteomic mass fraction, are potentially influenced by varying degrees of biodiversity within each taxonomic group. Yet, we restricted our analyses to these taxonomic groups due to the robustness of estimation with higher numbers of peptides (above simulations, and [43]). Further, this level of taxonomic resolution is typically used to compare ecological strategies across marine microbes, so we reasoned it would be useful to introduce these proteomic traits at the same level (e.g. [**?**]).

Here we outline the challenges in comparing taxonomic groups with varying biodiversity within each group, focusing on the environment-independent mass fraction proteomic ‘trait’. Biodiversity could influence the environment-independent mass fraction in several ways, depending on the exact meaning of ‘biodiversity’ in this context. The source of this variation could be due comparisons between taxonomic groups with varying levels of biodiversity (in terms of sequence diversity), or it could be due to a shift in community composition within taxonomic groups across samples (for example from one diatom species to another). These different mechanisms lead to different potential problems. For example, if community composition is constant across time, but one grouping is more biodiverse than another, our estimates could be interpreted as an average across subgroups (note this is not necessarily the case). But if there are significant shifts in community composition, then this might correspond with an apparent increase in peptide variability that arises from the change in community composition rather than changes in protein expression.

How could varying degrees of diversity be adjusted for? Simply correcting for total peptide diversity (in terms of numbers of peptides unique for a taxonomic group) is an obvious first step. Consider, however, the relationship between the total number of unique peptides for a species with a high regulatory cost (many regulatory proteins). There would be a causal connection between the number of unique peptides and the exact trait we are examining – regulatory cost – so ‘adjusting’ for peptide diversity would not be appropriate.

Another approach to assess variable biodiversity across taxa is to examine finer taxonomic resolution, and then estimate and compare the environment-independent proteomic mass fraction at that finer resolution with our original, coarse resolution estimates. This is problematic for two reasons: 1) peptides used for a finer taxonomic resolution are unlikely to be a random subsample, and certain protein functions are most likely enriched. If these protein functions are more or less likely to be constitutively expressed, estimates will not be comparable across taxonomic resolution would not be comparable. 2) Subsampling in mass spectrometry is explicitly biased towards highly abundant peptides. Peptides that are more abundant tend to have lower coefficients of variation (Supplementary Fig. S15). So, a subsample will systematically bias the environment-independent mass fraction upwards. We have outlined some of the principle challenges associated with using metaproteomics to estimate this proteomic trait, and future work is needed to address these issues. However, we think that this trait is still worth examining, because it likely underpins key aspects of ecological variability (e.g. as examined theoretically and experimentally in *E. coli*; [51]).

### 11.3 Abundance-Noise Relationship

Another potential bias in studying the environment-independent protein mass fraction is that less abundant proteins have more variation across identical conditions, as the mean protein coefficient of variation is negatively correlated with mean protein abundance in cultures [Supplementary Fig. S15 52]. So, identifying more peptides would increase the average coefficient of variation. However, we did not observe a negative correlation between the peptide-specific coefficient of variation and the mean peptide abundance (Supplementary Fig. S16), suggesting that this bias does not influence our estimated environment-independent peptide mass fraction.

**Figure S1:**
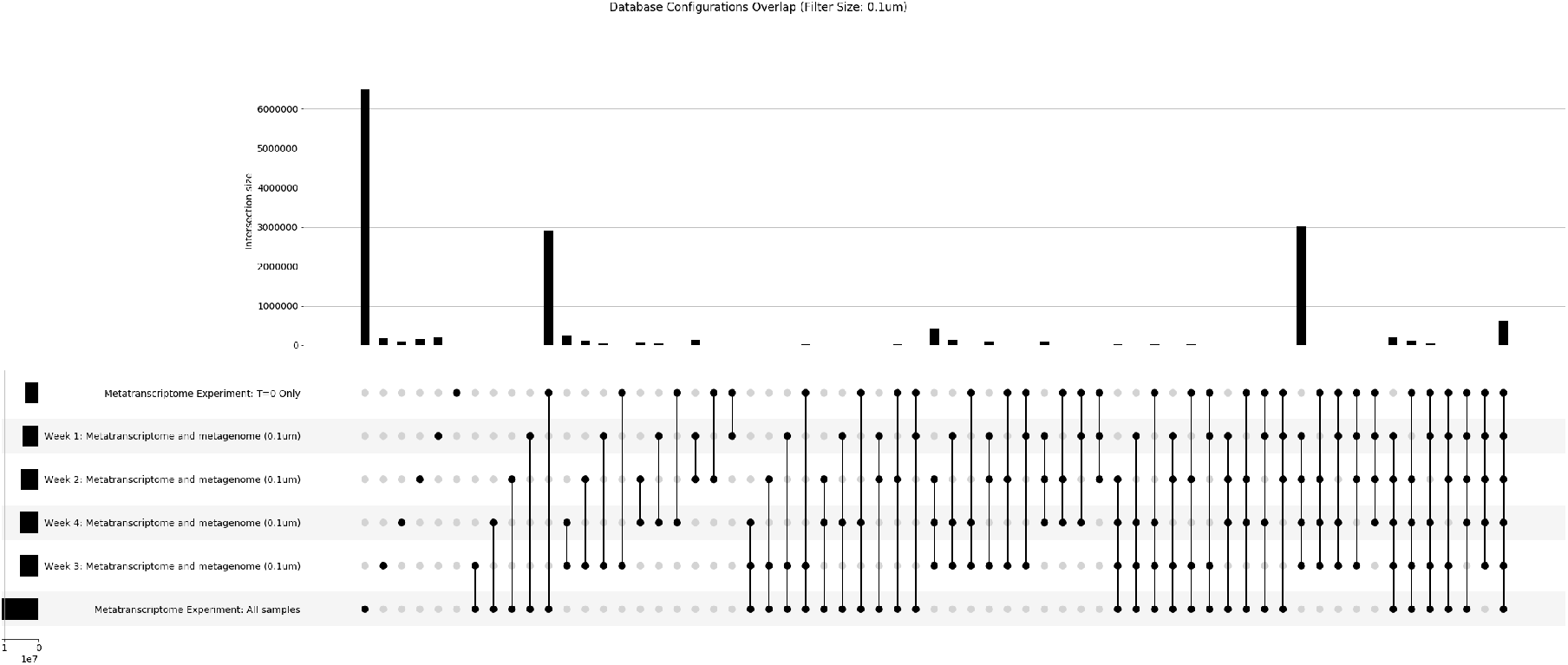
Representation of the overlap between different database configurations and the number of tryptic peptides within each. Bar graphs on top (with numbers above) represent the number of peptides identified with a given set of sequence groups (i.e. overlapping databases). The set of overlapping sequence groups is represented below with points and lines. The side bar plot, next to the database configuration name, is the total number of peptides within each sequence group database. In this figure, only the smallest filter size is shown (0.1 *μ*m).

**Figure S2:**
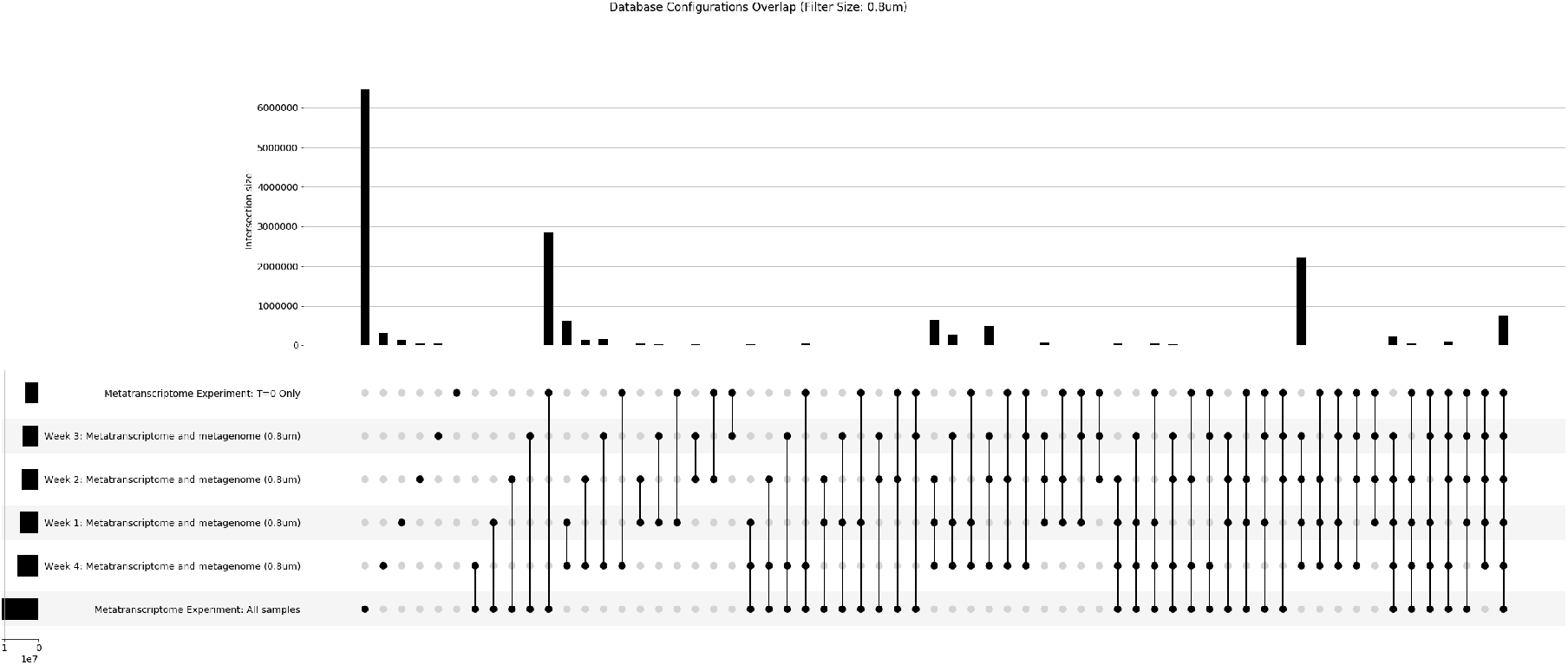
Representation of the overlap between different database configurations and the number of tryptic peptides within each. Bar graphs on top (with numbers above) represent the number of peptides identified with a given set of sequence groups (i.e. overlapping databases). The set of overlapping sequence groups is represented below with points and lines. The side bar plot, next to the database configuration name, is the total number of peptides within each sequence group database. In this figure, only the medium filter size is shown (0.8 *μ*m).

**Figure S3:**
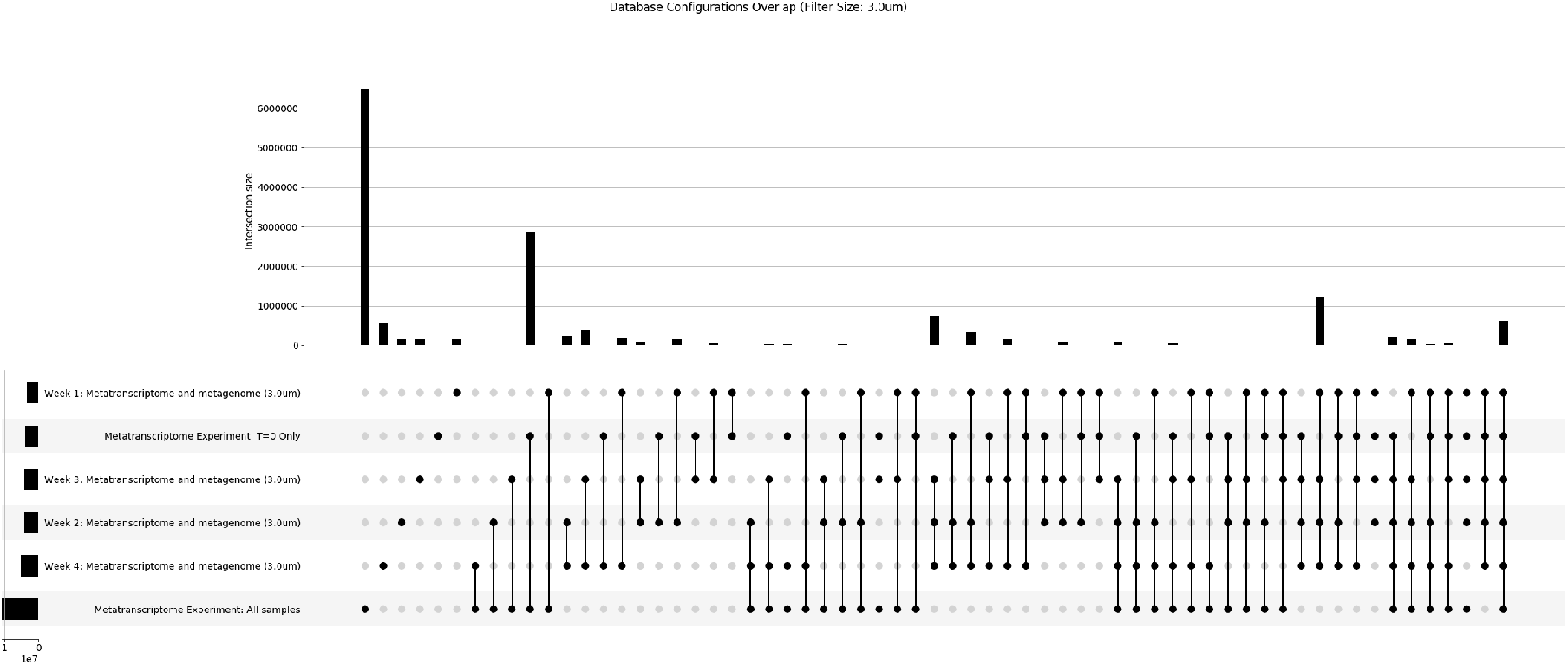
Representation of the overlap between different database configurations and the number of tryptic peptides within each. Bar graphs on top (with numbers above) represent the number of peptides identified with a given set of sequence groups (i.e. overlapping databases). The set of overlapping sequence groups is represented below with points and lines. The side bar plot, next to the database configuration name, is the total number of peptides within each sequence group database. In this figure, only the largest filter size is shown (3.0 *μ*m).

**Figure S4:**
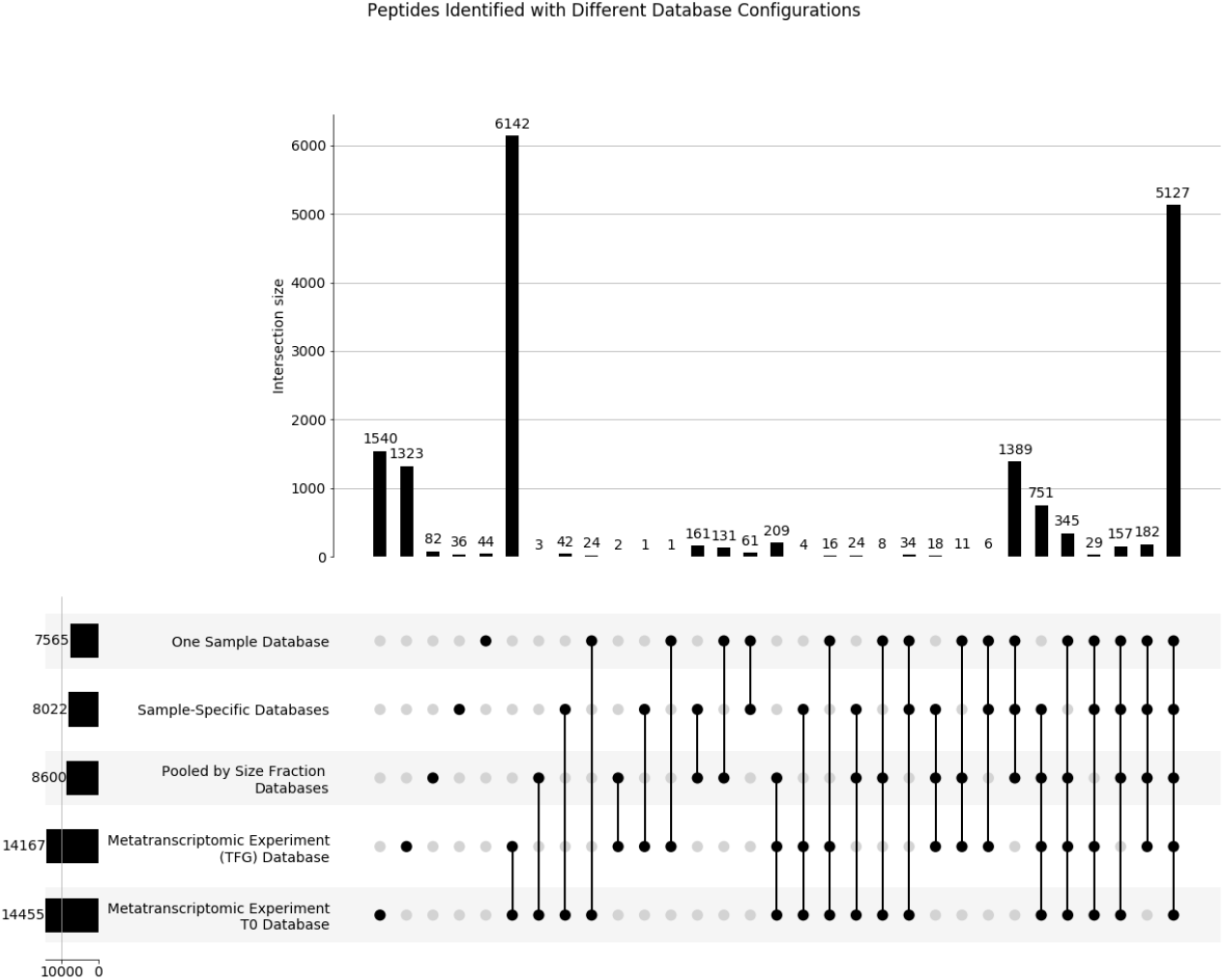
Representation of the overlap between different database configurations and the number of peptides identified with each. Bar graphs on top (with numbers above) represent the number of peptides identified with a given set of databases (i.e. overlapping databases). The set of overlapping databases is represented below with points and lines. For example, the first column on the left represents peptides uniquely identified using the database ‘Metatranscriptome Experiment T0’, where 1540 peptides were uniquely identified. The side bar plot, next to the database configuration name, is the total number of peptides identified using each database. In this figure, all filter sizes are summed together.

**Figure S5:**
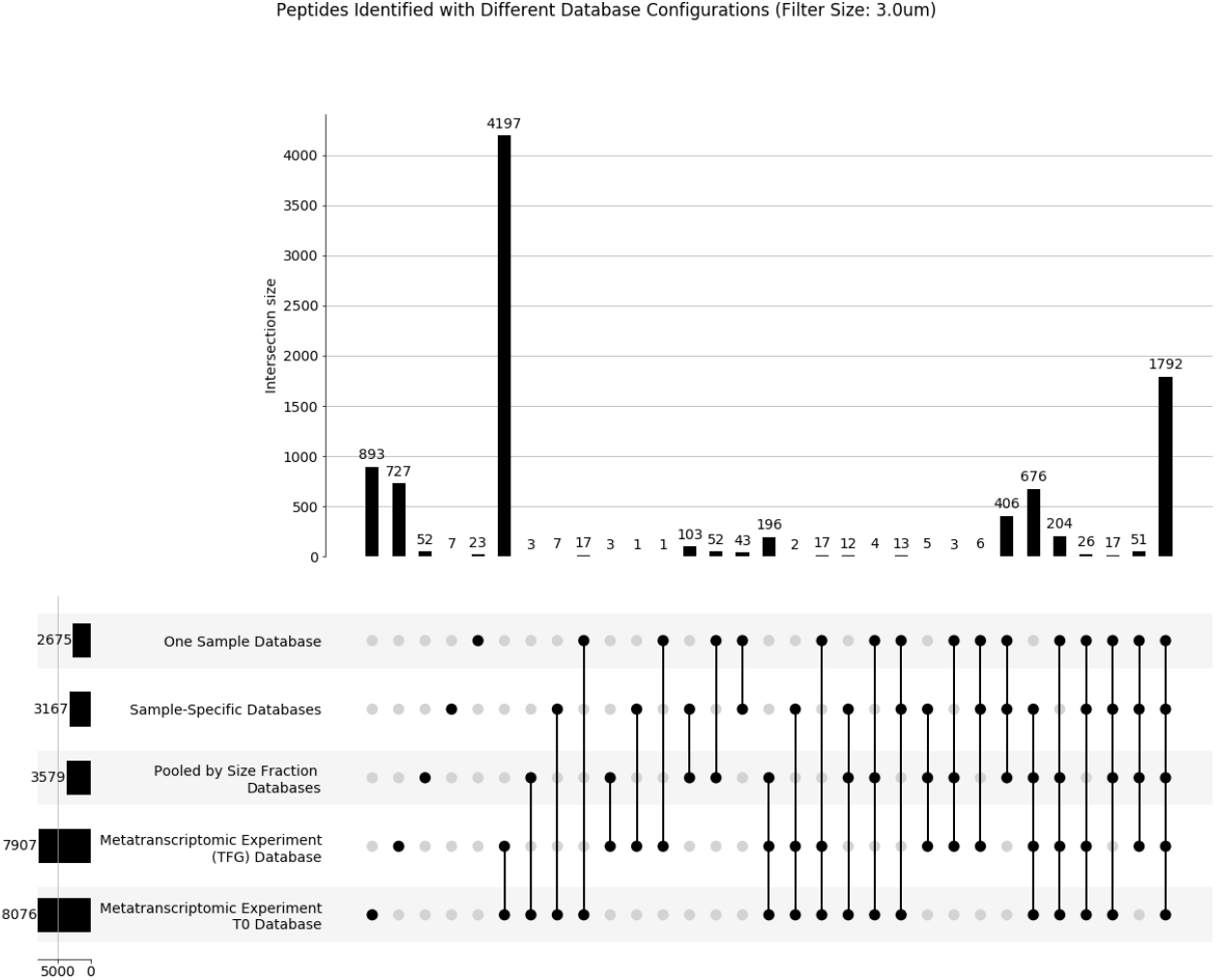
Representation of the overlap between different database configurations and the number of peptides identified with each. Bar graphs on top (with numbers above) represent the number of peptides identified with a given set of databases (i.e. overlapping databases). The set of overlapping databases is represented below with points and lines. The side bar plot represents the total number of peptides identified using each database. In this figure, only the largest filter size is shown (3.0 *μ*m).

**Figure S6:**
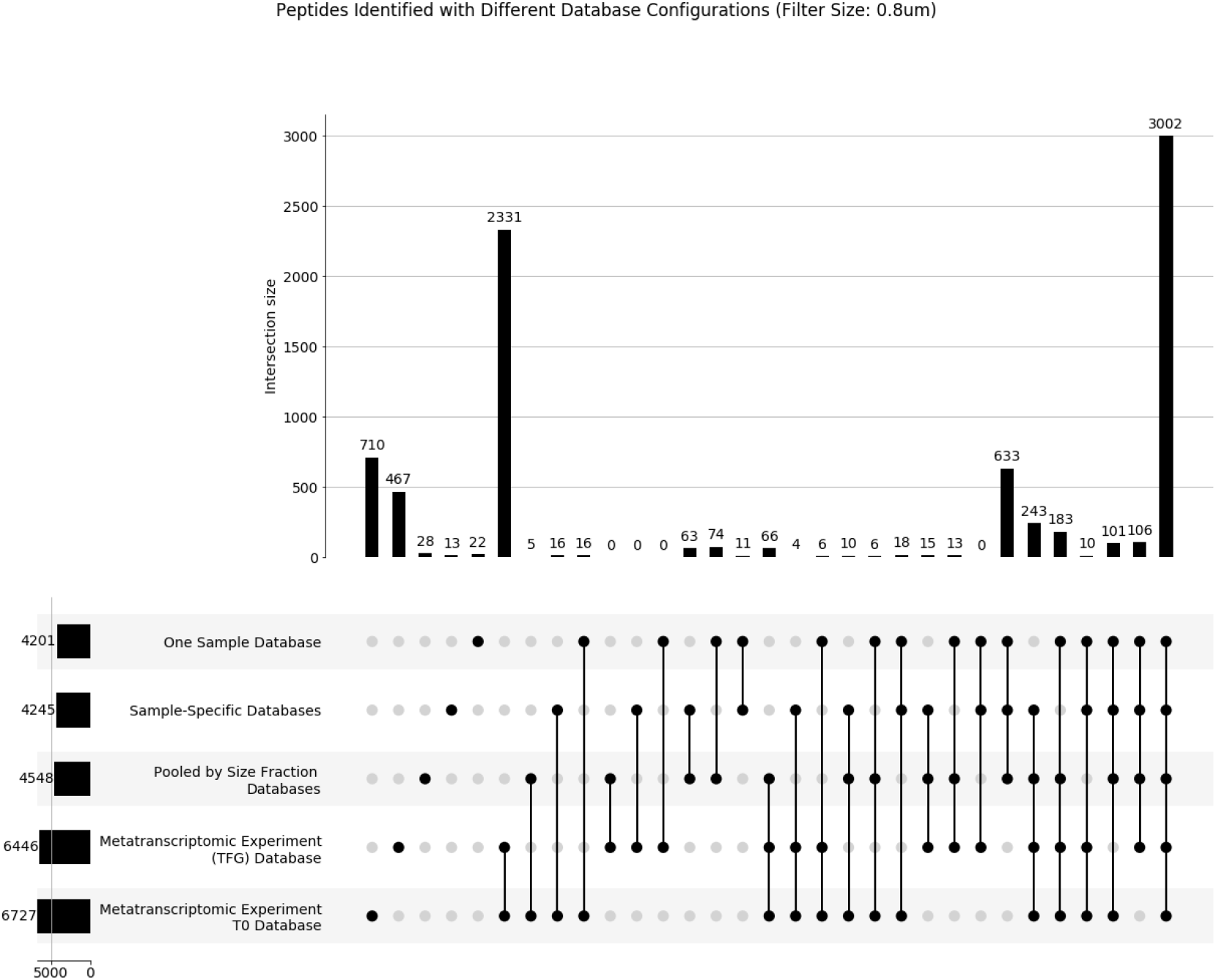
Representation of the overlap between different database configurations and the number of peptides identified with each. Bar graphs on top (with numbers above) represent the number of peptides identified with a given set of databases (i.e. overlapping databases). The set of overlapping databases is represented below with points and lines. The side bar plot represents the total number of peptides identified using each database. In this figure, only the middle filter size is shown (0.8 *μ*m).

**Figure S7:**
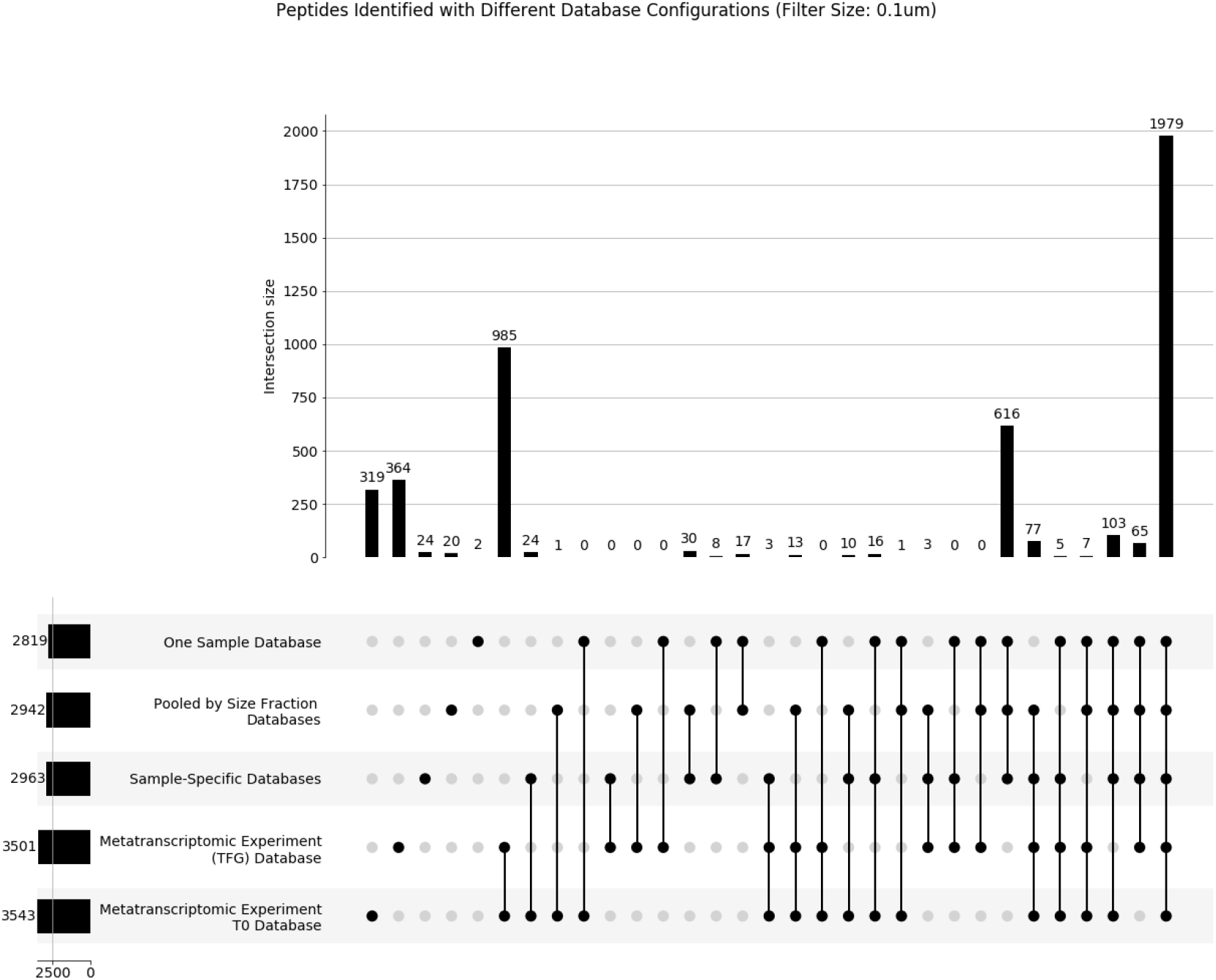
Representation of the overlap between different database configurations and the number of peptides identified with each. Bar graphs on top (with numbers above) represent the number of peptides identified with a given set of databases (i.e. overlapping databases). The set of overlapping databases is represented below with points and lines. The side bar plot represents the total number of peptides identified using each database. In this figure, only the smallest filter size is shown (0.1 *μ*m).

**Figure S8:**
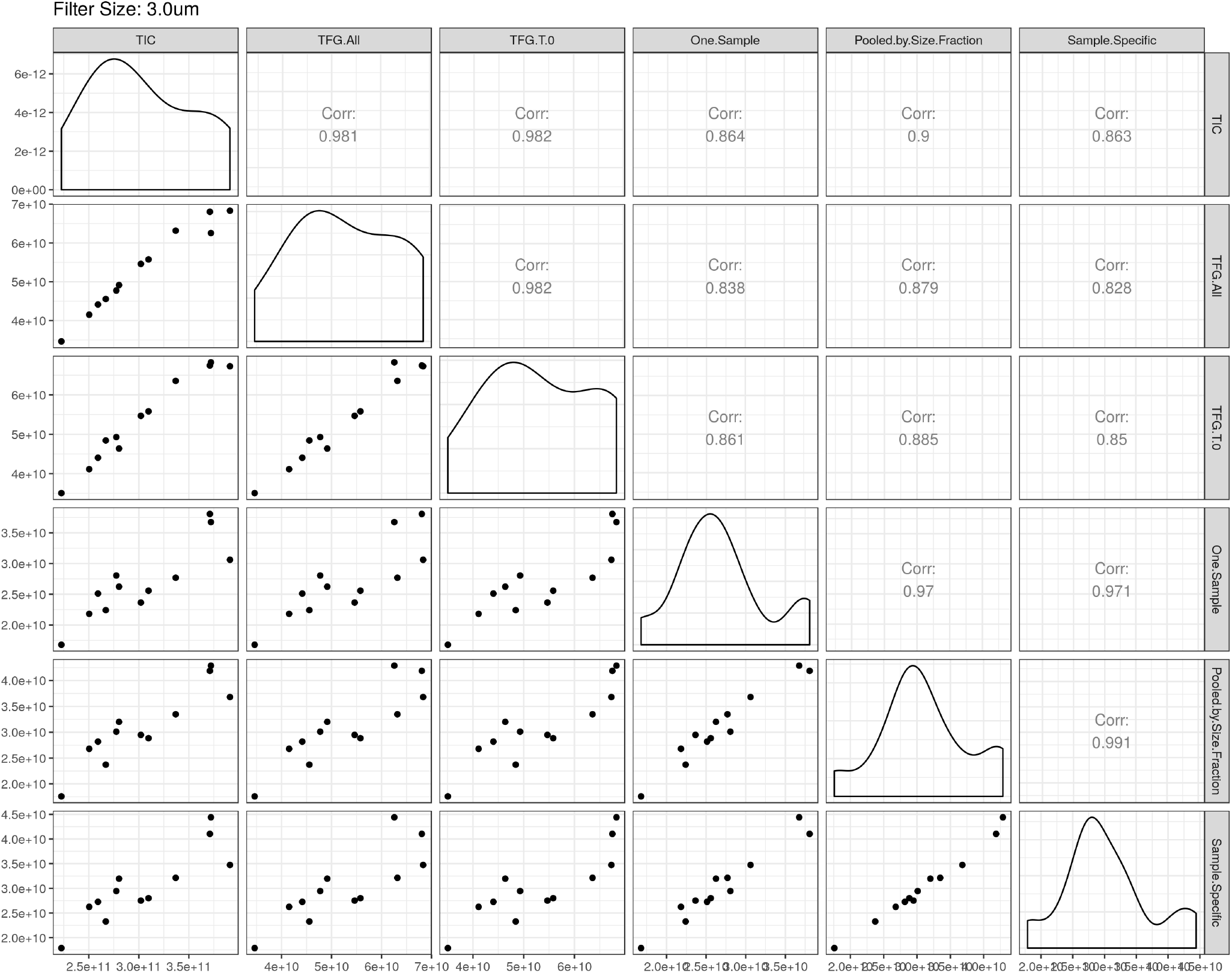
The sum of peptide intensities (i.e. normalization factors) are against each other for different database configurations, as well as against total ion current (TIC). Points represent different mass spectrometry experiments. Correlation values (coefficient of determination) are represented in corresponding locations. Only the largest filter size is shown here (3.0 *μ*m).

**Figure S9:**
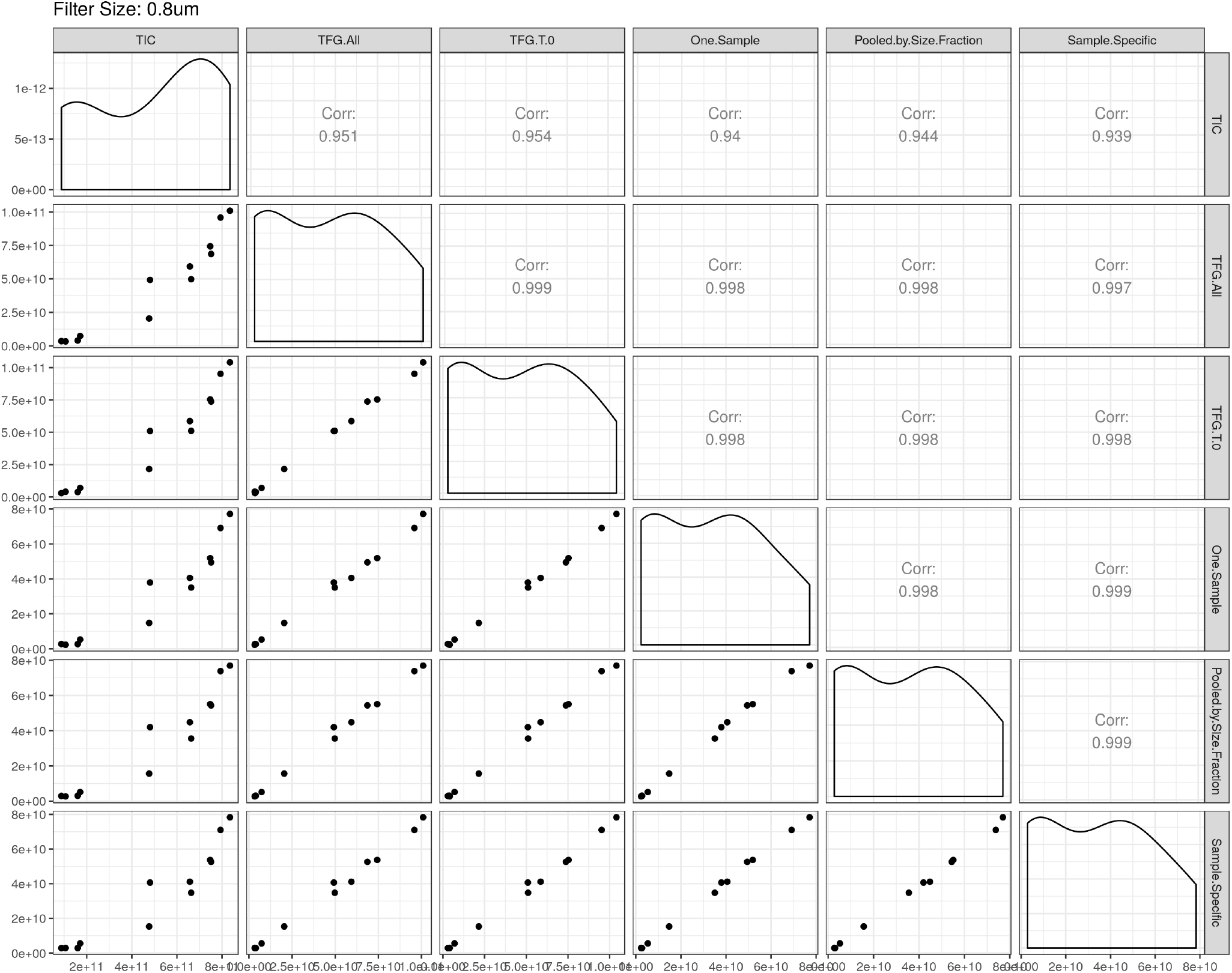
The sum of peptide intensities (i.e. normalization factors) are against each other for different database configurations, as well as against total ion current (TIC). Points represent different mass spectrometry experiments. Correlation values (coefficient of determination) are represented in corresponding locations. Only the middle filter size is represented here (0.8 *μ*m).

**Figure S10:**
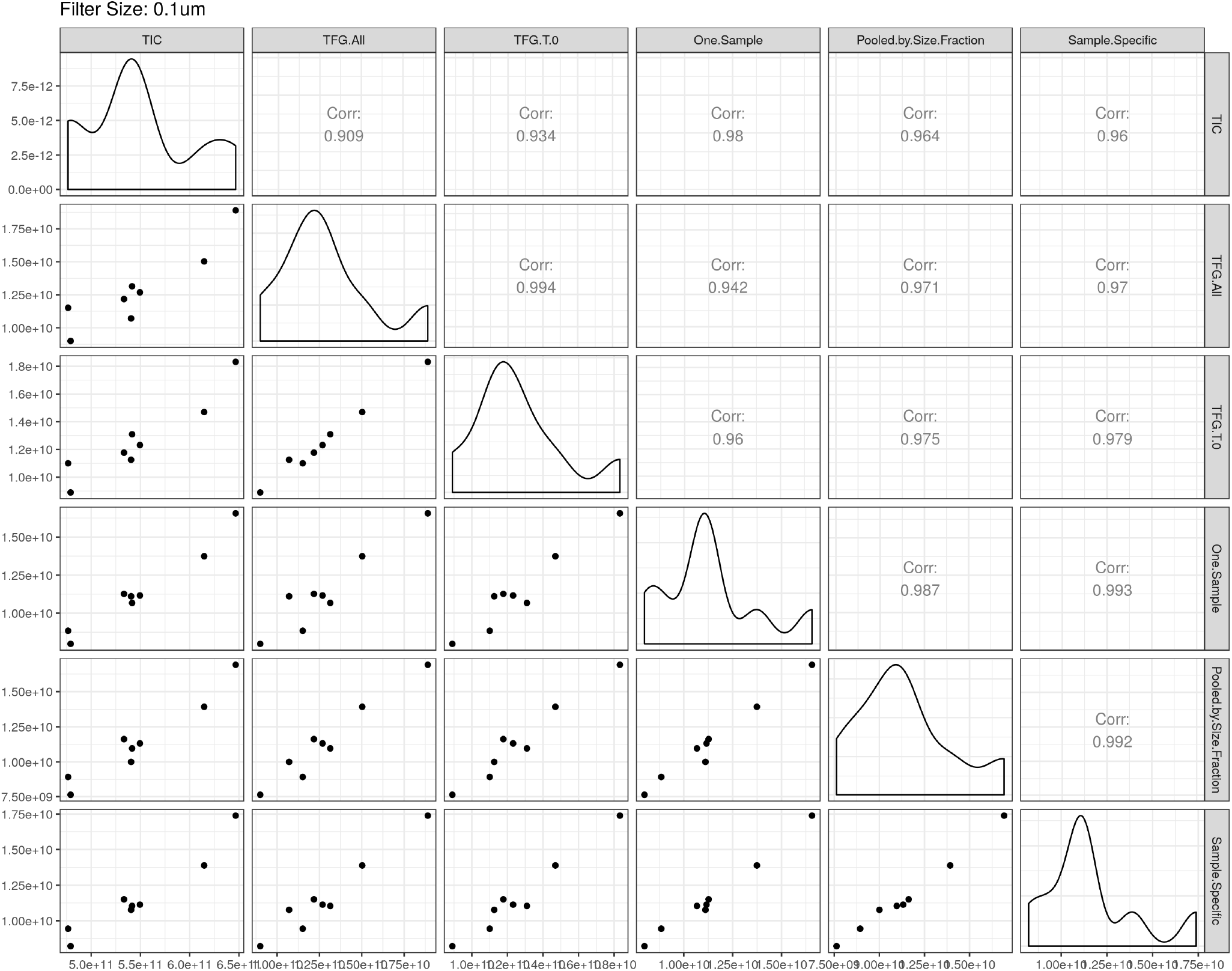
The sum of peptide intensities (i.e. normalization factors) are against each other for different database configurations, as well as against total ion current (TIC). Points represent different mass spectrometry experiments. Correlation values (coefficient of determination) are represented in corresponding locations. Only the smallest filter size is represented here (0.1 *μ*m).

**Figure S11:**
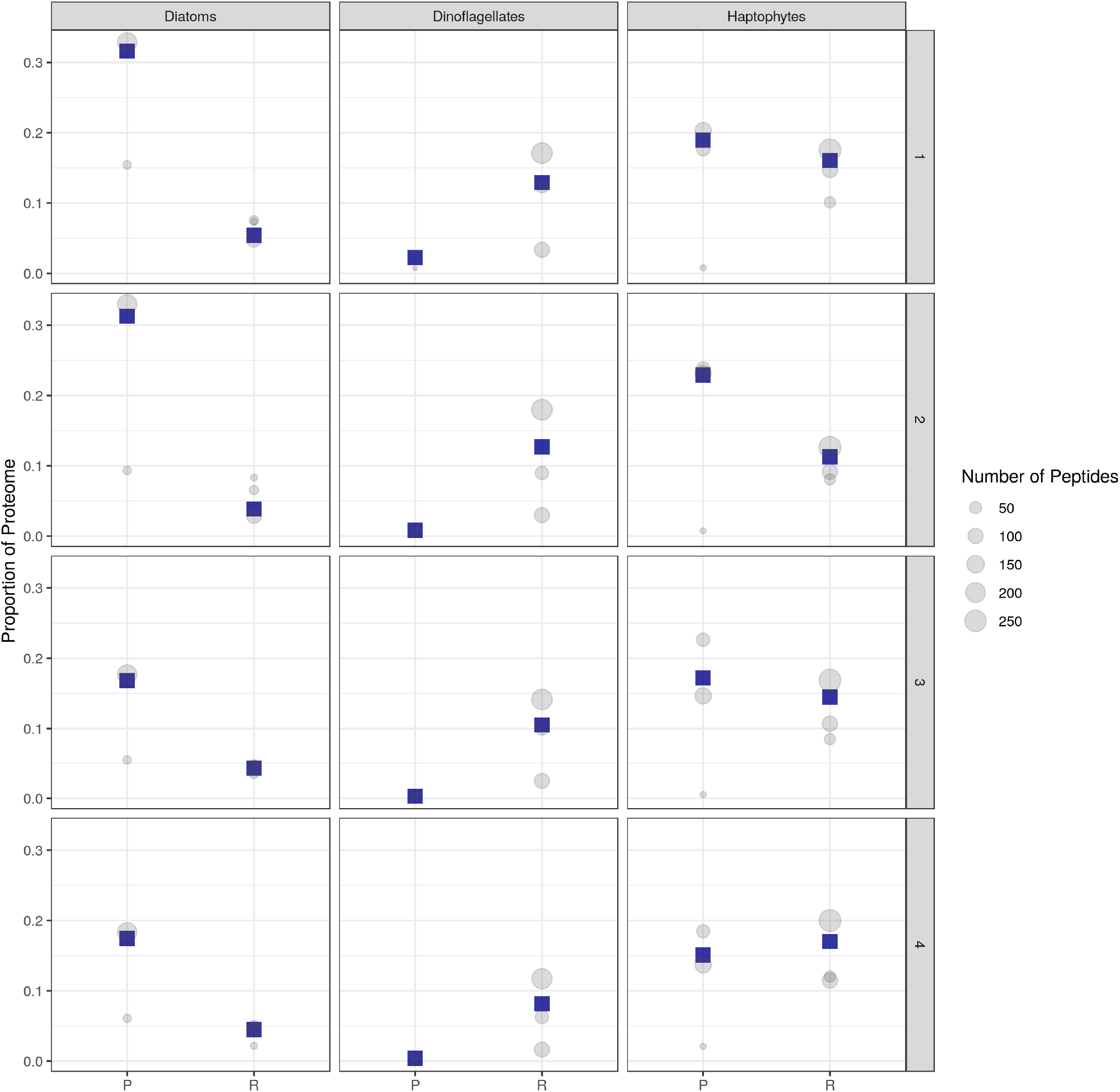
Demonstrating how various estimates across filters are collapsed into one estimate, based on the number of peptides identified within each filter size. Grey points represent the different filter sizes, while the size per grey point is the number of peptides observed in that filter, corresponding to a given taxa or a coarse-grained proteomic pool (P is photosynthetic protein pool, R is ribosomal protein pool). Dark blue squares are the weighted estimates. Numbers in the vertical direction (right side) correspond to the different sampling weeks.

**Figure S12:**
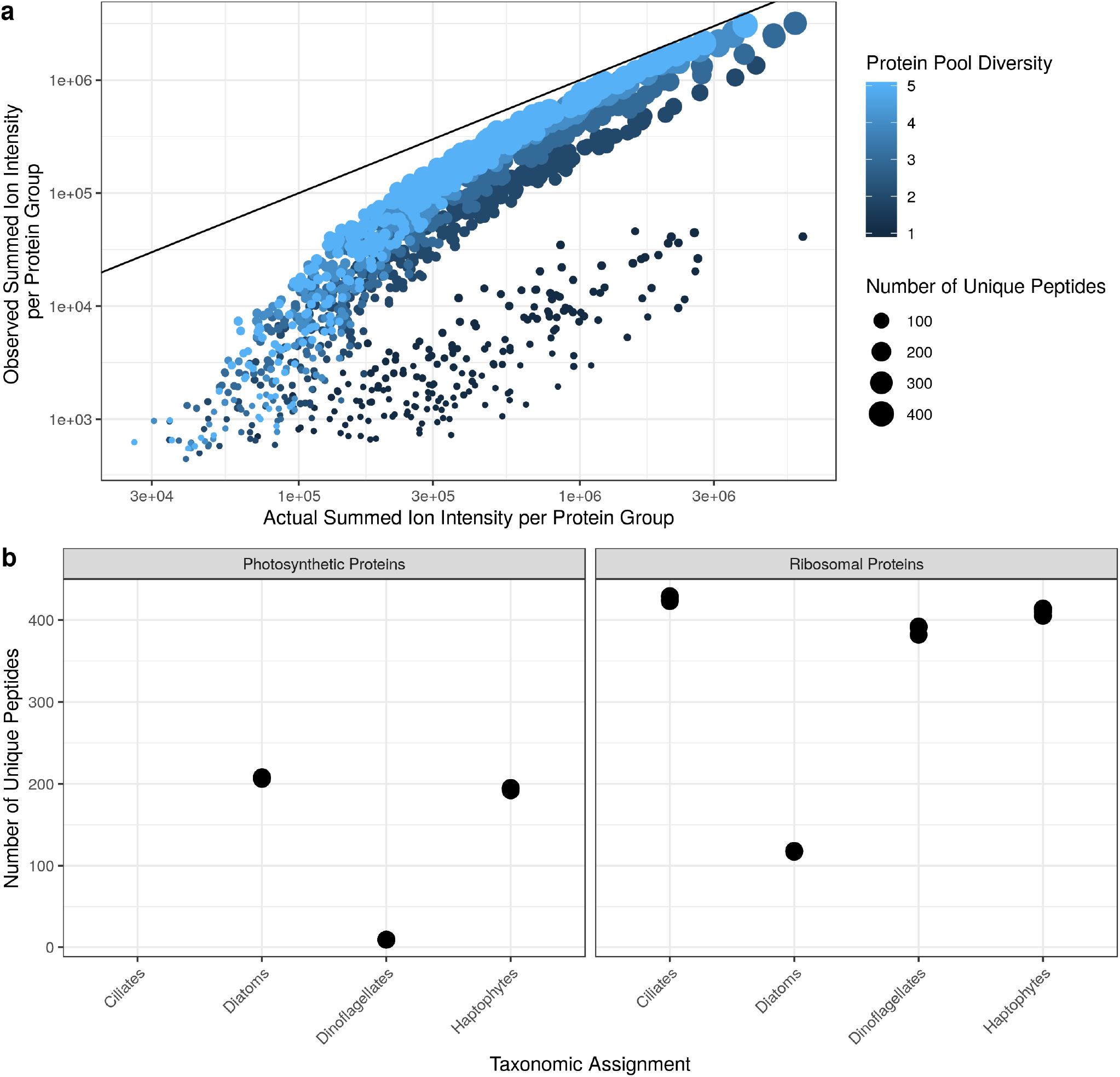
Results from the metaproteomic sampling simulation suggest that restricting analyses to protein groups and taxa that are abundant will prevent bias due to different amounts of diversity. **a**, varying degrees of simulated diversity (colour of points), demonstrates that low diversity pools are underestimated (i.e. far away from 1:1 line). Yet, if many peptides are observed (i.e. above 100 peptides), then the estimates are linearly correlated with the 1:1 line, with only slight underestimates due to diversity. **b**, the number of photosynthetic and ribosomal protein-specific peptides that are also taxon-specific is typically between 100–400 peptides. This suggests that our estimates of ribosomal and photosynthetic protein mass fraction not susceptible to this diversity-induced bias.

**Figure S13:**
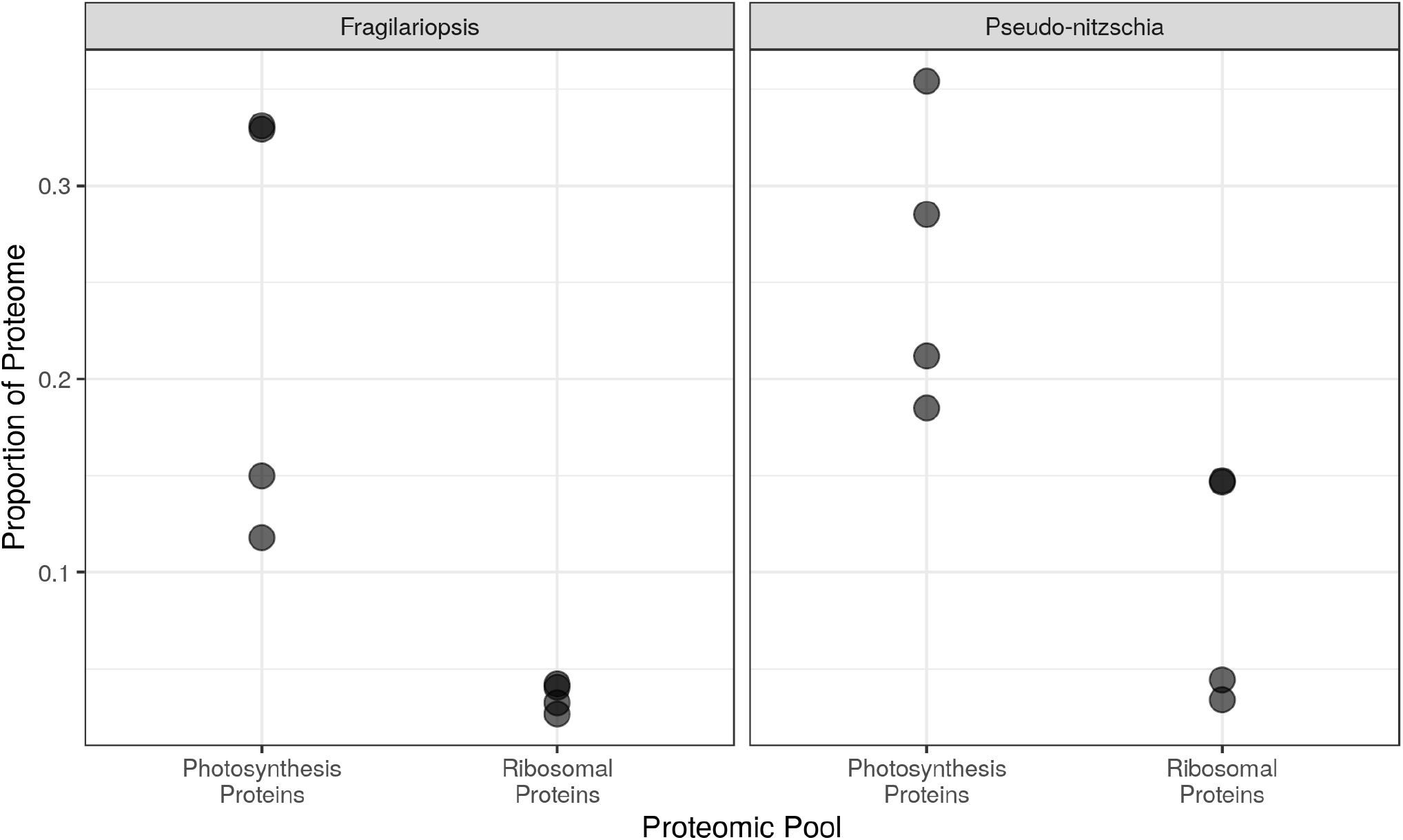
Proteomic proportions of two taxonomic groups of diatoms, *Fragilariopsis sp*. and *Pseudo-nitzschia sp*. Ribosomal and photosynthetic proportions were similar across groupings, and also similar to the larger grouping of diatoms.

**Figure S14:**
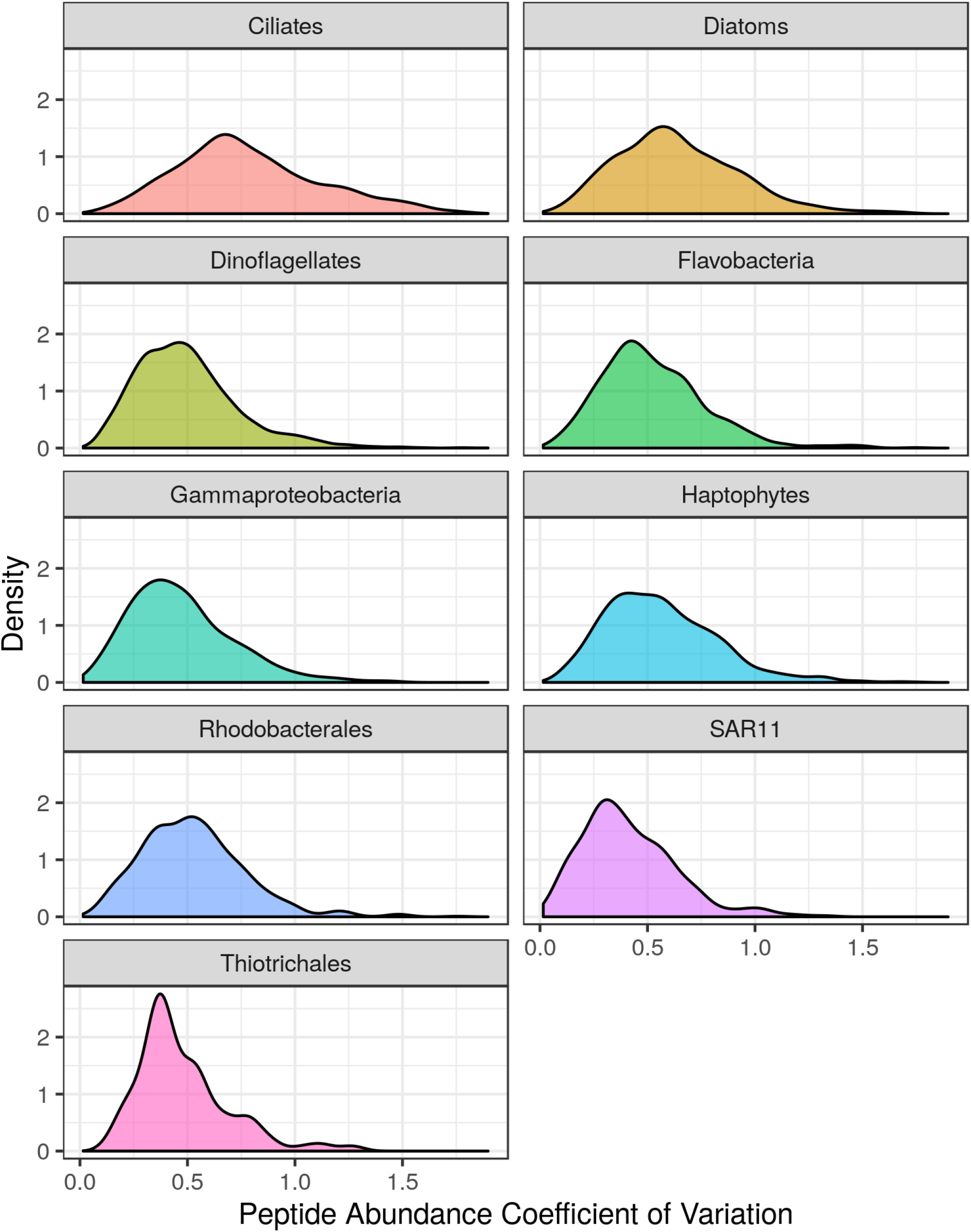
Distributions of peptide-specific coefficients of variation for each taxa we examined. In the main manuscript, only SAR11 and diatoms are shown. Methods for calculating this distribution are given in the main manuscript.

**Figure S15:**
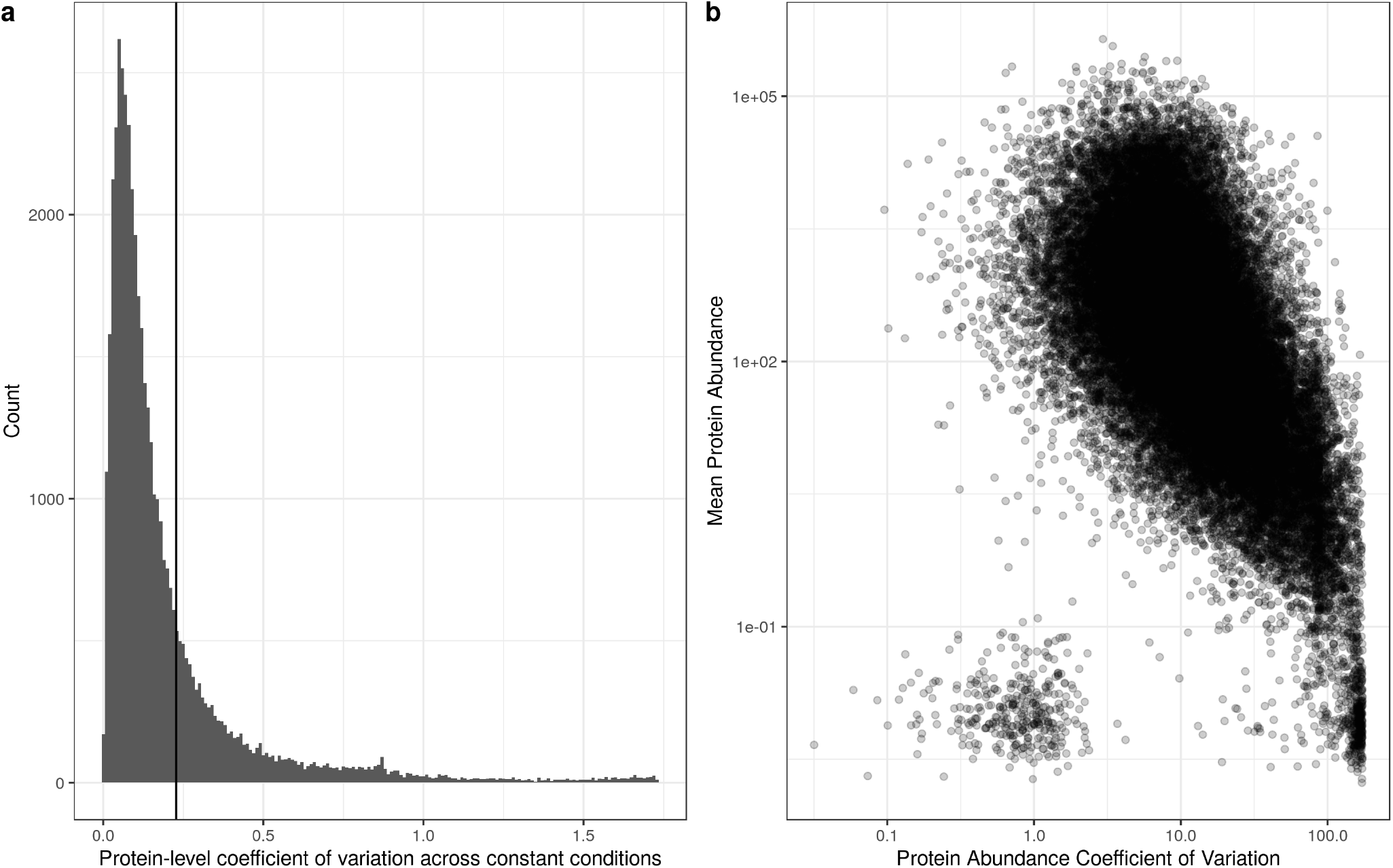
There is a negative correlation between the mean abundance value of a protein and its’ coefficient of variation.

**Figure S16:**
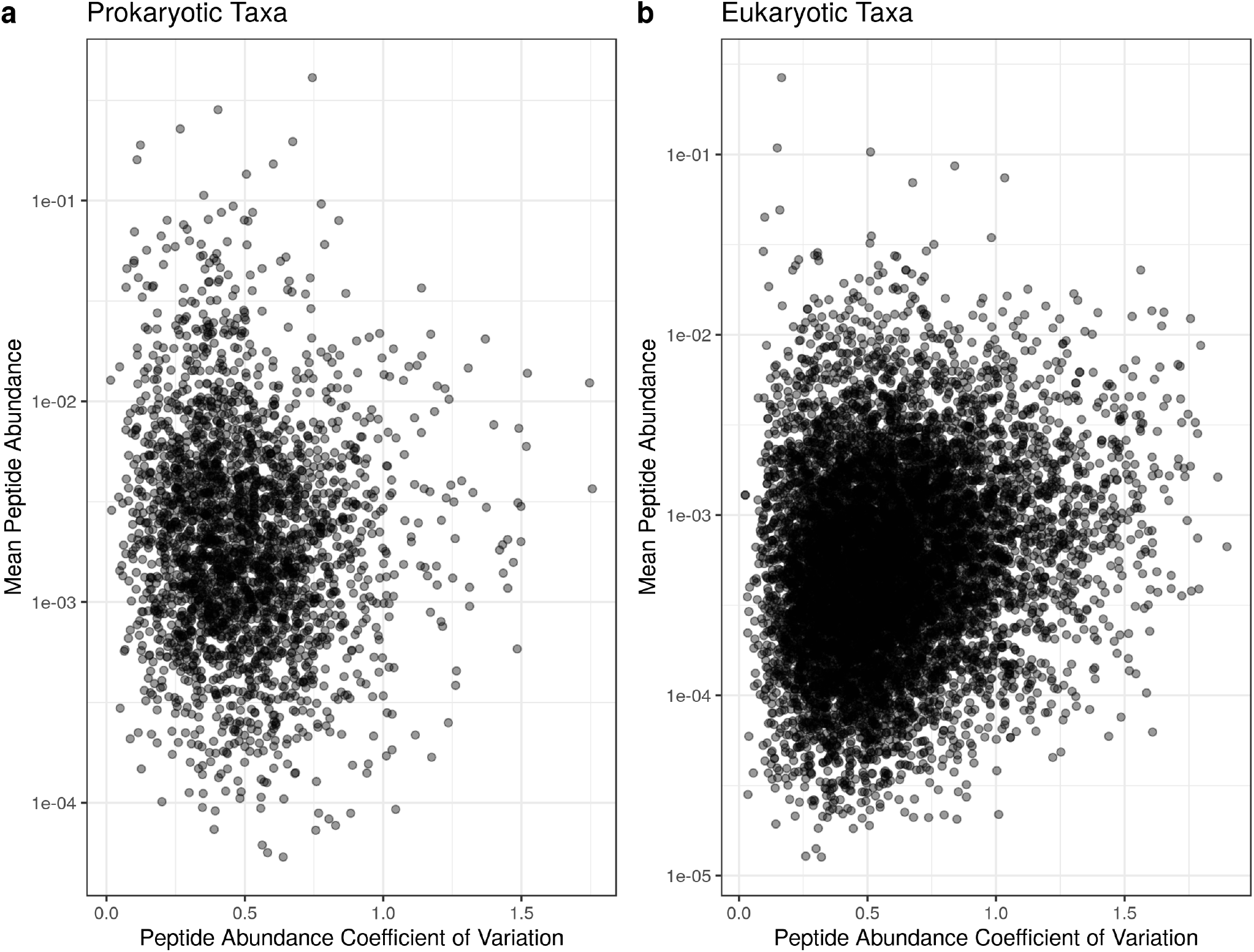
No significant relationship between the peptide abundance coefficient of variation and the mean peptide abundance for the prokaryotic and eukaryotic taxa we observed.

